# Potato Late Blight Control with a Botanical Product and Reduced Copper Applications

**DOI:** 10.1101/2025.01.08.627882

**Authors:** Tomke Musa, Andreas Kägi, Haruna Gütlin, Sylvain Schnee, Josep Massana-Codina, Karen E. Sullam

**Author notes:** Contributing authors.

## Abstract

Potato late blight (PLB), due to the pathogenic oomycete, *Phytophthora infestans*, can cause extensive economic damage, particularly in organic potato production. Although copper is used to combat PLB in organic production, it is banned in some countries, and its reduction or elimination as a plant protection product is an increasing priority in Europe. Alternative control strategies, including botanicals, could potentially reduce copper and control PLB. We investigated the application of *Frangula alnus* bark, its sequential use with a reduced copper application, and a reduced copper application alone in field and lab experiments. The influence of different dosages and preparations on efficacy and the quantity of the posited active ingredients were examined. *Frangula alnus* treatments decreased disease severity compared to a water control but showed differences in efficacy depending on dosage and disease pressure. Through in vitro and in planta experiments, we investigated whether *F. alnus* directly or indirectly controlled PLB. A bacterium (*Erwinia* spp.), originating from the *F. alnus* extract, colonized the media and accounted for most of the direct inhibition in vitro, but removing microorganisms through filtration had no effect on the extract’s efficacy in planta. The contribution of extract-associated microorganisms to PLB control is unclear and requires additional experimentation to assess. The presence of measured anthraquinones likely contributed to the effect of *F. alnus*. In field experiments, copper consistently and *F. alnus* generally (except for one year) reduced disease severity compared to a water control. No difference was observed in disease severity between the full and reduced copper treatments. Potato variety more consistently drove differences in total and marketable yields compared to the applied treatment. The relative stability of the yield suggests that treatment effectiveness is intertwined with the timing of disease development and environmental conditions.

## 1 Introduction

In many countries including Switzerland, organic potato production still relies on the use of copper to combat potato late blight (PLB) caused by the oomycete, *Phytophthora infestans* (La Torre et al, 2018). Copper, however, is a heavy metal that has long-term consequences for soil health and toxicity (Shabbir et al, 2020) and is already banned as a plant protection product by some countries, including Denmark, Estonia, Finland, the Netherlands, and Sweden (Tamm et al, 2022). In the last decades, copper reduction in organic potato production has become a priority (Tamm et al., 2022), and much research has been focused on alternative plant protection products, including the application of microorganisms or natural products ((Bangemann et al, 2014; Dorn et al, 2007). PLB is a polycyclic pathogen that typically requires multiple fungicide treatments within a growing season to control it, resulting in organic potato production receiving one of the highest fungicide and copper applications of all arable crops (Katsoulas et al, 2020; Andrivon and Savini, 2023). Therefore, the reduction of copper is tightly intertwined with the development of copper alternatives to control PLB.

Copper applications can also be reduced through the deployment of resistance varieties, but potato breeding requires a long-time horizon of at least 10 to 16 years (Keijzer et al, 2022; Pandey et al, 2023), which could be shortened to 7 to 13 years with the implementation of marker-assisted selection and genomic-estimated breeding values of complex traits (Slater et al, 2014). However, previous potato breeding efforts show that it is difficult to combine late blight resistance, tuber yield, starch content, and other characteristics (Reslow et al, 2022), and therefore, limited gains from breeding programs in Europe (Ortiz et al, 2022). Furthermore, the use of a single resistance gene is not a sustainable strategy, since P. infestans strains can evolve to overcome qualitative resistance (Gilroy et al, 2011). Genome-editing techniques could facilitate the development of resistant varieties, but the public, particularly organic producers and consumers, do not accept genetically modified organisms’ use in potato breeding (Pacifico and Paris, 2016). Therefore, adopting lower copper application levels and pairing them with an alternative product can help reduce copper application levels on a shorter-time scale. Previous experiments with botanicals demonstrate efficacy against oomycetes (Forrer et al, 2017; Mulholland et al, 2017; Taillis et al, 2022; Dagostin et al, 2010), and buckthorn alder (*Frangula alnus*) bark extract from material that has been dried and finely ground is among the botanicals that show potential to combat potato late blight (Forrer et al, 2017). However, further optimizing the preparation and dosage are needed to enable the economic feasibility of the treatment.

The goal of this work was to determine the efficacy of the extract of dried and ground *F. alnus* bark, gain information about its accumulative effects, evaluate its sequential use with reduced copper treatments, and optimize its usage. In order to determine the efficacy of different *Frangula alnus* preparations in the field, we carried out field experiments over four growing seasons to test various preparations and dosages. In addition, we conducted lab experiments, including in planta and in vitro assays and analytical chemistry to gain more information about the nature of the active compounds, treatment dosage and timing, and applicability to practice. Overall, this work highlights the possibility of further reducing copper allowances and the application of sequentially used treatments.

## 2 Methods

### 2.1 Field experiment set-up

From 2019 to 2022, four potato field experiments were conducted at Agroscope in Zurich Reckenholz, Switzerland. A split plot design was used in the experimental fields with six replications in 2019 and 2020 and five replications in 2021 and 2022. Each plot included two sub-plots: one sub-plot with two rows of the variety Agria and another sub-plot with two rows of the variety Victoria. Agria and Victoria are common varieties used in Swiss potato production, including organic production (Swisspatat, 2023). These varieties are classified as moderately susceptible to late blight according to the recommended Swiss Variety List of potatoes (Schwärzel et al, 2019). Varieties with moderate susceptibility are recommended for the evaluation of low-risk products as they are expected to be less effective than higher-risk products (Evenhuis, 2021). Each sub-plot was 7.5 m^2^ and contained two rows that were each 75 cm wide and 5 m long. In each row, the 15 planted tubers were spaced 33 cm apart. The placement of Agria and Victoria subplots within the plots was randomly assigned. In the four rows between the plots, two rows of a low susceptible variety (Panda in 2019 and 2020, Jelly in 2021, and Innovator in 2022) were planted along the long plot edge. The highly susceptible variety, Bintje, was planted between the two rows of the low susceptible variety (see Supplementary Figure A1 for more information on field layout) to act as spreader rows. The low susceptible variety was changed in different years due to availability. Treatments were randomly assigned to plots within each block.

For all field experiments, certified seeds were used, and the fields were managed according to the European and Mediterranean Plant Protection Organization guidelines (EPPO, 2021) for conducting fungicide efficiency experiments against *P. infestans* on potatoes. Briefly, it provides standards on how to conduct trials to evaluate fungicide efficacy against *P. infestans*, including trial conditions, layout, treatment application and disease assessment. The only deviations from these guidelines were regarding plot length and variety choice. Although a longer plot (8 m) is prescribed for yield evaluations, the shorter acceptable plot length (5 m) was used because of field size constraints. Additionally, varieties used in this study are considered moderately susceptible to PLB to better mimic conditions in organic potato production citepSchwaerzel. In every year except 2022, PLB occurred naturally in the field experiments. In 2022, untreated leaves from the susceptible cultivar Bintje that had been infected naturally in a nearby untreated field were laid in the Bintje spreader rows to promote the infection within the experiment. Haulms of Bintje spreader rows were destroyed when 5-15% of the leaves were diseased. More details on the experimental set-up and agronomic conditions can be found in the supplementary material (Supplementary Tables A1, A2, and Figure A2).

#### 2.1.1 Treatments

All experiments contained a water control, a copper full dose (0.4 kg/ha) treatment, a copper reduced dose (0.2 kg/ha) treatment, and one or more *F. alnus* treatments consisting of different dosages or preparations 1. The dosage of copper was chosen based on a schedule with 10 treatments that result in a total application of 4 kg/ha per year, the maximum copper amount allowed in Switzerland. The number of treatments increased across the years as more information about the dosages and extraction methods became available through laboratory experiments. Starting from the field experiment’s second season, treatments with a reduced *F. alnus* dosage were added. Additionally, an *F. alnus* treatment with a lyophilization step during its extraction (see section below for details on preparation) was included in the third and fourth season of the field experiment. For copper treatments, Kocide Opti^®^ (Bayer Agrar Schweiz AG, Basel, Switzerland) was used, which contains copper hydroxide in the form of a water dispersible granule. The water control (treatment 1) was sprayed with the same amount of water as the treated plots during the applications. Treatment number 9 consisted of a water treatment for the first four applications before switching to the reduced copper dose, and it served as a control for treatment number 8. Treatment 8 included four initial *F. alnus* treatments followed by six copper treatments.

#### 2.1.2 Preparation of *Frangula alnus* solution

For treatments 4, 5 and 6 (Table 1), dried and milled bark of *F. alnus* (Faulbaumrinde PhEur, Dixa AG, St. Gallen, Switzerland, country of origin Poland) was suspended in tap water and stirred at room temperature (see Supplementary Table A2 for water volumes). In 2019 and 2020, the *F. alnus* solution was stirred for 2 hours as previously published (Forrer et al, 2017). The *F. alnus* treatments used 2021 and 2022 were prepared with a reduced stirring time of 30 minutes because a laboratory experiment revealed that the efficacy was unchanged between stirring times (see section below on extraction procedure experiments for details). After stirring, the solution was filtered through a 0.2 mm cheesecloth and then used within two hours for the treatments in the field. For treatment 7, the extraction method of *F. alnus* was modified according to a previously described procedure (Massana Codina, 2020). In short, powder-ground bark of *Frangula alnus* was extracted at 20% w/v using aqueous extraction in nanopure water by stirring for three hours at room temperature. The resulting extract was then centrifuged (4000 rpm, 10 min, 20 °C) and the supernatant was filtered through a ø 9 cm 589/3 cellulose filter (Schleicher & Schuell, Dassel, Germany) using a vacuum. The aqueous solution was then freeze-dried. This modified procedure will be referred to as the low dose, freeze-dried (FD) treatment (Table 1). The fine powder obtained was stored in a sealed box at 10 °C in darkness. Before field applications, the amount of the fine powder that corresponds to the original bark weight of the low dose treatment (i.e. 20% of the original weight) was suspended in tap water and stirred for 30 minutes in the same manner as the *F. alnus* treatments. All treatments were then directly used in the field following their preparation.

**Table 1:** Treatments and dosages applied for potato field experiments conducted in the years 2019 - 2022. Dosages below show amount sprayed (per ha and treatment), and these dosages were applied for all 10 sprayings. Equal dosages of active ingredients from the treatments were applied during each spraying. Each treatment was mixed with tap water based on a total spray volume of 450 L/ha for the first four applications. To account for the growing canopy, the amount of water was increased for subsequent applications to 700 L/ha in 2019-2021 and 630 L/ha in 2022. The full dosage of the copper treatment 0.4 kg/ha applied 10 times equals the maximum allowed dosage in Switzerland.

| Years | Treatment Number | Treatment | Amount per ha |
| --- | --- | --- | --- |
| 2019-2022 | 1 | Untreated control (water) | NA |
| 2019-2022 | 2 | Copper full dose | 0.4 kg<sup>1</sup> |
| 2019-2022 | 3 | Copper reduced dose | 0.2 kg |
| 2019-2021 | 4 | *Frangula alnus* full dose | 40 kg<sup>2</sup> |
| 2020-2022 | 5 | *Frangula alnus* medium dose | 25 kg |
| 2021-2022 | 6 | *Frangula alnus* low dose | 2.5 kg |
| 2021-2022 | 7 | *Frangula alnus* low dose (FD) | 0.5 kg<sup>3</sup> |
| 2021-2022 | 8 | 4x *Frangula alnus* low dose, followed by 6x reduced dose copper | 2.5 kg (*F. alnus*)/0.2 kg (copper) |
| 2022 | 9 | 4x no treatment (water), followed by 6x reduced dose copper | 0.2 kg |

<sup>1</sup>The copper weight refers to the copper amount in the product Kocide Opti<sup>®</sup>, which is 30% copper w/w in the form of copper hydroxide.
<sup>2</sup>The *Frangula alnus* weight refers to the dried ground bark used to make the extract.
<sup>3</sup>The 0.5 kg of extracted freeze-dried material is the product of 2.5 kg of *F. alnus* ground bark that was used as a starting material. Therefore, the amount of original material is the same as the low dose treatment (treatment 6).

#### 2.1.3 Field experiment spray application

Dosages during applications were consistent throughout the season (Table 1), although spraying volume increased to accommodate the growing plants (Supplementary Table A2). The first application was carried out according to the recommendation of the Swiss decision support system PhytoPRE (www.phytopre.ch) (Cao et al, 1997; Steenblock and Forrer, 2002). The system considers late blight occurrence (date, location and distance), growth stage of the crop as well as weather conditions favorable for the development of *P. infestans*. All subsequent applications were performed in a weekly routine and were generally applied in the morning unless rain postponed the spraying times. The applications were shifted according to the weather forecast to try to avoid application on days with rainfall, and treatments were aimed to be applied one day in advance when rain was forecasted (see Supplementary Figure A2 for meteorological conditions during each season and on treatment days). For some applications, spraying on days with rainfall was unavoidable. For those occurrences, it was attempted to postpone the treatment until after precipitation or ensured that the spray coating could dry before rainfall. A total of ten applications was made each year (see Supplementary Table A2).

In 2019 and 2020, the applications were conducted with a plot sprayer (Schachtner Gerätetechnik, Ludwigsburg, Germany) equipped with a 1.5 m spray-boom and three IDK compact air induction nozzles (IDK yellow, IDK 90 -02 C, Lechler GmbH, Metzingen, Germany) at 2.2 bar. In 2021 and 2022, the treatments were applied with a tractor (Fendt® F 345 GT, AGCO GmbH) with a 3 m spray-boom and 6 IDK nozzles (IDK red, IDK 120-04 C). For all years, a spraying volume of 450 L/ha was used for the first four applications. To cover the entire plant canopy, the 6 top-down nozzles were complemented with 5 dropleg nozzles (Lechler tongue nozzles, FT 140-02) as soon as the plants reached BBCH 31-35 corresponding to a switch to the larger spraying volume (630 L/ha in every year except for 2022 when 700 L/ha was used). In 2020, because of extremely wet soil conditions, the fifth treatment was applied with a knapsack sprayer (Birchmeier, Stetten, Switzerland) outfitted with a 3 m spray- boom and 6 IDK nozzles (IDK brown, IDK 120-05 C, 1.9 bar). The spraying pressure was adjusted according to the equipment used.

#### 2.1.4 Data collection from field experiments

The infection severity was estimated based on the area of infected plant tissue within each variety sub-plot (Forbes et al, 2014). In June and the beginning of July, more frequent observations were performed (2 times per week) to better assess the early development of the disease, while later in the season (mid and late July), the assessments were made weekly. Potatoes were harvested and yield was calculated from the two internal rows of each plot. The potatoes were sorted into four size classes (<42 mm, 42-55 mm, 55-70 mm, >70 mm). The middle two size classes (42-55 mm and 55-70 mm) constitute the marketable potatoes and were weighed to calculate the marketable yield. During the harvest, the tubers were examined for tuber blight. Additionally, 40-50 tubers per treatment were then examined for signs of tuber blight one month after harvest. For plots in which fewer than 50 tubers were harvested, all tubers were examined (min. per sub-plot was 40 tubers).

### 2.2 Climate chamber and laboratory experiments

Laboratory experiments were concurrently conducted with field experiment to better understand the extraction procedure, dosages, spraying time and mechanism of *P. infestans* control with *F. alnus*. These experiments consisted of detached leaf assays and mycelium growth experiments as well as the chemical analyses of the *F. alnus* solutions.

#### 2.2.1 Extraction procedure and application timing experiment on detached leaves

Detached leaves of the susceptible variety Bintje were used to compare the efficacy of *F. alnus* after 30 versus 120 minutes of extraction. A 4% concentration of *F. alnus* was prepared in the same manner as the field experiment explained above. The aqueous solution of *F. alnus* was stirred for 30 minutes in addition to the treatment stirred for 120 minutes.

Within this experiment, the timing of the *F. alnus*application in relation to *P. infestans*inoculation was also tested. The three treatment levels consisted of applications either: 1) one day before *P. infestans* inoculation 2) two days before *P. infestans* inoculation, or 3) both two days and one day before *P. infestans* inoculation. A control treatment composed of deionized water was included, and it was sprayed according to the same regime. A gravity feed spray gun with cup (DeVilbiss, England) was used to apply 3 mL of the treatments to each terminal leaf at a pressure of 1 bar. All treatments included eight detached leaves that were each placed separately on a 7 x 7 cm square of wire mesh that had been laid over a moistened filter paper in individual round dishes with a diameter of 125 mm and height of 34 mm (Semadeni AG, Switzerland). The dishes were closed with a lid and were set at a slight angle to allow for the water to collect at the bottom, and they were randomly distributed in a blocked design within a climate chamber. The experiment was run at 18 °C and a 16-8 hr day-night schedule of which 2 hours consisted of a gradual reduction or increase of light. During the night period, the humidity was 85%, and 75% during the day.

A *P. infestans* polyspore isolate (polyspore isolate no: 18-001) collected in 2018 from untreated Bintje leaves in Reckenholz, Zurich was used for this and all laboratory and climate chamber experiments presented in this study. Its SSR genotype (Martin et al, 2019) had been categorized to “Other” (David Cooke, *Pers. comm.*), and it is an A1 mating type. The sporangia inoculation solution prepared by scraping mycelia from a two-week old rye agar plate and diluting the sample to a sporangia concentration of 5 x 10^5^/mL. The spore preparation was incubated at 4 °C for two hours in the dark prior to inoculation to promote the release of zoospores. Each detached leaf consisted of the terminal leaflet and two first primary leaflets. The terminal leaflets were inoculated with *P. infestans* two times while each primary leaflet was inoculated once, totaling four inoculation points across the three leaflets (See supplementary Figure A3). A volume of 30 *µ*L of the *P. infestans* sporangia preparation was used for each inoculation point. The humidity within the climate chamber was increased to 100% for 48 hours following the *P. infestans* inoculation, during which the climate chamber was kept dark. After the 48 hour period, the previous day-night (16-8 hr) schedule and humidity settings (day: 85% and night: 75%) were reinstated. The percentage of leaf area that was diseased with *P. infestans* was visually assessed and estimated seven days post infection.

#### 2.2.2 *Frangula alnus* dosage experiment

Detached leaf assays were carried out in the same manner as described above to test different application amounts of *F. alnus* and to determine its half-maximal inhibitory concentration (IC50). Previous studies had used 4% *F. alnus* extracts, which corresponded to approximately 40 kg/ha (Forrer et al, 2017; Krebs et al, 2006). In order to obtain a more economically feasible *F. alnus* treatment, reduced *F. alnus* treatment amounts were tested in laboratory experiments to assess the efficacy of lower application amounts in a controlled environment. A five-point dilution series of *F. alnus* was used for the treatment amounts, ranging from 250 kg/ha to 0.025 kg/ha. To connect the laboratory dosages to relevant values for the field, we assumed that there are 40,000 plants/ha and each plant has 55 compound leaves. Therefore, for the highest dose, 250 kg/ha, a preparation of 11.364 g of *F. alnus* dried and ground bark was mixed with 300 mL of tap water. From this solution, a serial dilution was made for each dosage. For this experiment, detached leaves of the variety Bintje were used. To obtain the treatment with the highest concentration, a 4% *F. alnus* bark was extracted in tap water with stirring for 30 min and then strained with a 0.2 mm cheesecloth. From this concentration, the subsequent four treatments were serially diluted using tap water. The detached leaves were sprayed as described above with 3 mL of the treatment per leaf at 1 bar until the treatment was well distributed on both sides of the leaf. For the negative control, tap water was applied directly to the detached leaves.

Each treatment included 12 detached leaves that were randomly distributed according to a block design within the climate chamber. The climate chamber conditions and *P. infestans* strain used were the same as those used for the extraction procedure and spray timing experiment. The leaves were sprayed two days prior to artificial infection with *P. infestans*, which was applied as 4 drops of 30 *µ*L of sporangia solution, including two drops on the terminal leaf and one drop on each of the first two primary leaflets. The sporangia preparation was prepared in the same manner as in the experiment described above, and the sporangia concentration used was 1 x 10^5^/mL. The disease severity was visually assessed by estimating the surface area of *P. infestans* infection on each detached leaf at seven days post infection. The IC50 was calculated from this experiment using the 4-parameter logistic regression model from the XLSTAT software (Lumivero). For the model, disease severity data was the dependent variables, and the extract concentration was the explanatory variable.

#### 2.2.3 *In vitro* mycelial growth experiment

In this experiment, *F. alnus*’s direct control on *P. infestans* mycelial growth was tested in vitro applying the same treatments used in the field (treatment numbers 1-7, Table 1) on rye agar (200 g untreated rye seed boiled and strained with a normal kitchen strainer, 5 g D-glucose, 20 g agar in 1000 mL water). *Frangula alnus* powder was prepared in the same manner as described above with a 30 min extraction step, but the tap water was autoclaved prior to the aqueous extraction step. Due to observations that a characteristic bacterial culture would grow on rye agar from *F. alnus* extractions, each of the *F. alnus* treatments were applied with and without an additional filtration step to eliminate the direct influence of microorganisms associated with the plant extracts. The bacterium was later identified to be an *Erwinia* spp. from sequencing the 16S rRNA gene (GenBank accession PV124383, see Supplementary Material for details). Subsequent experiments revealed that the *Erwinia* bacterium could be cultured from multiple batches of *F. alnus*. Three batches were tested, and all three yielded an *Erwinia* spp. bacterium (see Supplementary Material for additional information). The extent of the bacterium’s co-occurrence with *F. alnus* bark is not known.

For the filtered solutions, two supplementary filtration steps were made that included 1) a vacuum filtration step using a paper filter (Schleicher & Schuell AG, 8714 Feldbach ZH) and 2) a manual filtration step using a 20 mL Luer-Lok syringe (BD Plastipak, Becton Dickinson GmbH, Germany) equipped with a 0.2 *µ*m filter (Minisart®, Sartorius stedim biotech, Germany). Following filtration, the solution was plated to verify that the bacteria that grew on non-filtered *F. alnus* had been removed. A ø 5 mm *P. infestans* mycelial plug from a freshly cultivated colony (within 10-14 days) was placed upside down in the center of each Petri dish (ø 9 cm, Greiner Bio-one, Switzerland). Six concentric wells were constructed 15 mm from the dishes’ center and were filled with 25 *µ*L of the test substance, of which only one was used on each Petri dish. Each test substance was repeated in 8 dishes. Any dishes that showed contamination were later excluded from the analysis except for the *Erwinia*spp. that grew in the treatments of the non-filtered aqueous extractions of *F. alnus*. A second iteration was run with a subset of the treatments, including the negative control (autoclaved deionized water), Copper (full), Frangula low, Frangula low (filtered). After seven days, the mycelial growth of *P. infestans* from the original plug was quantified as the percentage of mycelial coverage on each dish. For the measurements, the whole plate area was quantified with Image J (Schneider et al, 2012), and a modified ImageJ macro (Laflamme et al, 2016) was used to measure the area of a defined color threshold in pixels (See Supplementary Material). The experiments were normalized based on a comparison between each treatment’s mycelial growth to its negative water control within each experiment and then combined for statistical analysis. The difference between the mycelial coverage in the treatment and the negative water control was used to determine the treatments’ growth inhibition.

#### 2.2.4 Detached leaf assay with filtered and non-filtered *Frangula alnus* extracts

Because bacterial growth was observed following plating *F. alnus* solutions on rye agar (see the in vitro experiment described above and Supplemental Material for more information), detached leaf assays were run to determine whether filtered and non-filtered *F. alnus* extracts differed in their efficacy in controlling *P. infestans*. Only the high dosage of *F. alnus* from the field experiment was included in the experiment with a positive control (copper) and negative control (autoclaved deionized water). A filtered bacterial free extraction of high dosage *F. alnus* was prepared as described in the in vitro experiment, and the same protocols for the detached leaf set-up was followed as described in the detached leaf experiments focusing on dosage and *F. alnus* extraction and application timing. The assay was conducted two times with replicates of ten leaves per treatment in each assay.

Leaves of the variety Bintje were placed individually in a Petri dish (125 mm, Semadeni AG, Switzerland) on a 7×7 plastic mesh lined with paper towels, and 8 mL of tap water was added to the bottom of each Petri dish to keep the leaves fresh. The upper side of the leaf was sprayed with 1.5 mL of the prepared treatments at 1 bar and incubated in a climate chamber at 18 °C with 14 h day light and 75% humidity followed by 10 h darkness and 85% humidity. One day after treatment application, the leaves were infected at four points with a 30 *µ*L *P. infestans* sporangia suspension. The spore suspension was prepared as described above with a final concentration of 1.60 x 10^5^ sporangia per mL for the first assay (Assay A) and 1.75 x 10^5^ sporangia per mL for the second assay (Assay B). One week post inoculation, the percentage of the infected leaf area was visually assessed as described above.

### 2.3 Chemical profiling and anthraquinone quantification from *Frangula alnus* bark

In order to investigate the chemical profile and determine the putative active compounds that may be involved in the *F. alnus* extracts’ control of *P. infestans* in the field, freshly prepared extracts used in the field experiments were immediately frozen (within 30 minutes) at -20 °C for later analysis. In total, five preparations of the medium dosage, low dosage and low-FD dosage from two applications in 2021 and three applications in 2022 were analyzed. Additionally, the preparations from the aqueous extraction time experiment were profiled.

All preparations were thawed, filtered through a 22µm filter and directly injected on a Vanquish Horizon UHPLC System consisting of a degasser, a mixing pump, an autosampler, a diode array detector (PDA) and a Vanquish Charged Aerosol detector (ThermoFisher, Waltham, MA, USA). Samples were separated on an Atlantis BEH C18 column (2.1 x 100 mm, 1.7µm, Waters, Milford, MA, USA) using a mobile phase consisting of nanopure water containing 0.1% formic acid (A) and acetonitrile containing 0.1% formic acid (B); to obtain a chromatographic profile of the extract, a separation was performed using a step gradient of 5% to 40% B in 30 minutes, from 40% to 100% in 5 minutes, and isocratic at 100% for 3 minutes at a flow rate of 0.3 ml/min. In order to quantify frangulins and emodin, the separation was performed using a step gradient from 15% to 30% B in 4 minutes, isocratic at 30% B for 13 minutes, from 40 to 100% in 10 minutes and held at 100% B for 4 minutes at a flow rate of 0.3 ml/min. Detection was performed at 210, 254, 280, 350 and 435 nm, and the CAD conditions were the following: evaporator temperature set at 35 °C and the Data Collection Rate set at 10 Hz.

### 2.4 Statistical analysis

All data analyses were conducted in R (version 4.4.0) (R Core Team, 2023). Area under the disease progress curve (AUDPC) was calculated using the R package agricolae (de Mendiburu, 2023). For field experiments, disease severity based on the AUDPC and marketable and total yield were compared separately for each year using linear mixed models with the experimental blocks as a random factor and variety, treatment, and their interaction as fixed factors using the R package nlme (Pinheiro et al, 2023). The residual values were checked for normality and homoscedasticity using a Q-Q and scale-location plots, respectively. To better fit the assumption of normality, the yield data were logarithmically transformed. Pairwise comparisons between treatments were performed using the estimated marginal means, and Tukey’s honest significant difference tests were used to adjust p-values in the emmeans package (Lenth et al, 2024). The compact letter display function (cld) in R package multcomp was used to display differences between groups (Hothorn et al, 2008). The measurements of compounds extracted from the F. alnus preparations were compared using linear mixed models as above, and these data were logarithmically transformed prior to analysis to account for the deviation from normality.

All data analyses were conducted in R (version 4.4.0) (R Core Team, 2023). Area under the disease progress curve (AUDPC) was calculated using the R package agricolae (de Mendiburu, 2023). For field experiments, disease severity based on the AUDPC and marketable yields were compared separately for each year using linear mixed models with the experimental blocks as a random factor and variety, treatment, and their interaction as fixed factors using the R package nlme (Pinheiro et al, 2023). The residual values were checked for normality using a Q-Q plot. Pairwise comparisons between treatments were performed using the estimated marginal means, and Tukey’s honest significant difference tests were used to adjust p-values in the emmeans package (Lenth et al, 2024). The compact letter display function (cld) in R package multcomp was used to display differences between groups (Hothorn et al, 2008). The measurements of compounds extracted from the *F. alnus* preparations were compared using linear mixed models as above. However, the data were logarithmically transformed prior to analysis to account for the deviation from the model’s assumptions of homoscedascity and normality.

The climate chamber experiment on extraction duration and spraying regime was analyzed with Kruskal-Wallis Rank Sum Test to explore differences between *F. alnus* and control treatments as well as pair-wise differences within the experiment. In order to assess the effects of the *F. alnus*’s extraction time, treatment application regime, and their interaction, they were included in a zero-inflated negative binomial model using the glmmTMB package (Brooks et al, 2017) with experimental blocks as random factors. All other laboratory experiments, including the detached leaf tests focusing on effects of *F. alnus* dosage and *F. alnus* filtration as well as the mycelia growth inhibition tests were analyzed with Kruskal-Wallis tests, followed by Conover’s all-pairs rank comparison tests using the R package PMCMRplus (Pohlert, 2018) with the Benjamini & Hochberg adjustment for multiple comparisons (Benjamini and Hochberg, 1995).

## 3 Results

### 3.0.1 Field Experiments

The 2021 potato growing season was the most conducive for disease development followed by 2020. Accordingly, disease severity was highest in 2021, followed by 2020 and 2019, and in 2022, there was very little PLB (Figure 1). In all years, the treatment had a significant effect on the AUDPC, while variety had a significant effect on the AUDPC in the years 2019 and 2021, and no interaction between the treatments and variety was found in any of the years (Table 2). The *F. alnus* treatments showed significantly reduced AUDPCs compared to the water control (Figure 2) except in 2021 when high amounts of precipitation led to increased disease pressure (Supplementary Figure A2, Supplementary Table A1). In the field, the reduced copper was found not to be significantly different from the full copper treatment. The marketable and total yields showed significant variety effects in all four years but only significant treatment effects in 2019 and 2020. Compared to the copper treatments, the Victoria plots treated with the high dosage of *F. alnus* showed a trend of reduced yields in 2019 and 2020, which was found to be significant in marketable yield in 2019 and total yield in 2020 (Figure 3). The cultivar, Agria, had no yield reduction nor did the medium dosage when applied in 2020. In 2021 and 2022, the yields were not different among the treatments within each variety. Treatment and variety interactions did not have a significant effect on either marketable or total yield (Table 2).

**Fig. 1:**
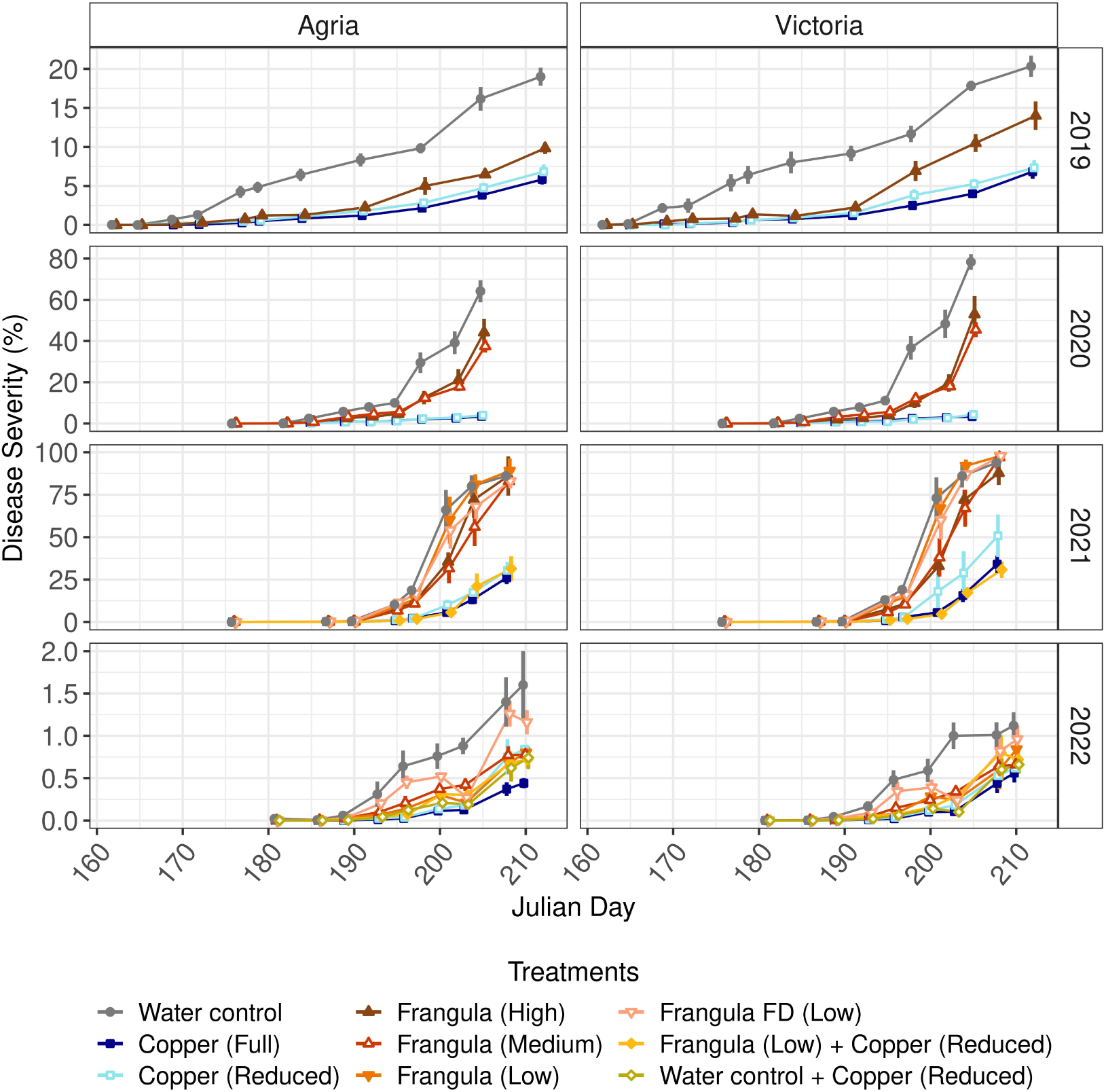
Disease severity of potato late blight epidemics in two varieties from field experiments between 2019 and 2022. Treatments are described in Table 1. The y-axis scale changes according to the year, showing the high variation in disease severity across years. Bars indicate standard error of the means.

**Fig. 2:**
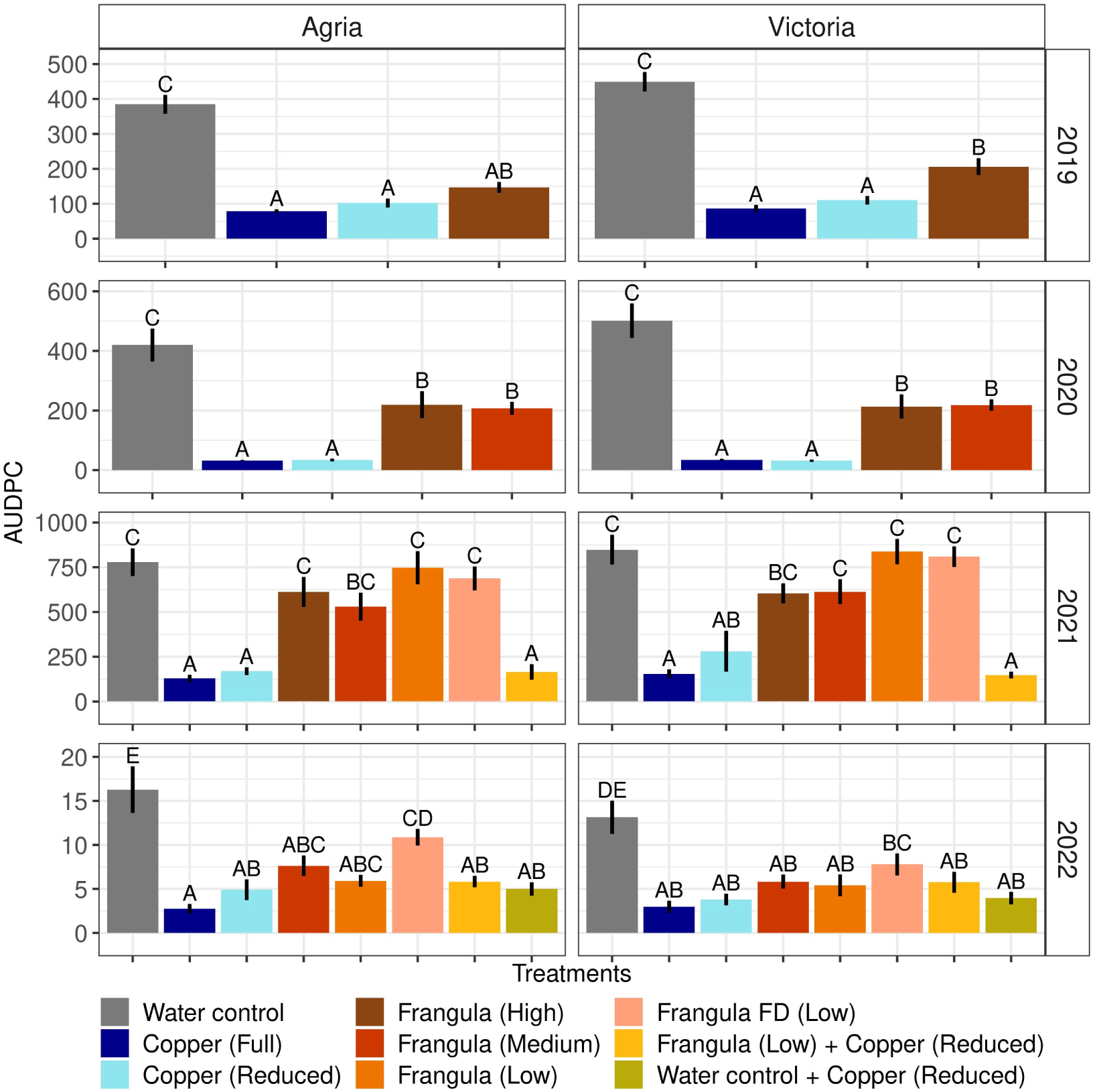
Area under the disease progress curve (AUDPC) of potato late blight in two varieties from field experiments between 2019 and 2022, showing treatment by color as described in Table 1. The scale of the y-axis varies according to the year, reflecting the substantial variation in disease severity, particularly from 2021 to 2022. Treatments that do not share grouping letters were shown to be significantly different from each other. Bars indicate the standard error of the mean.

**Fig. 3:**
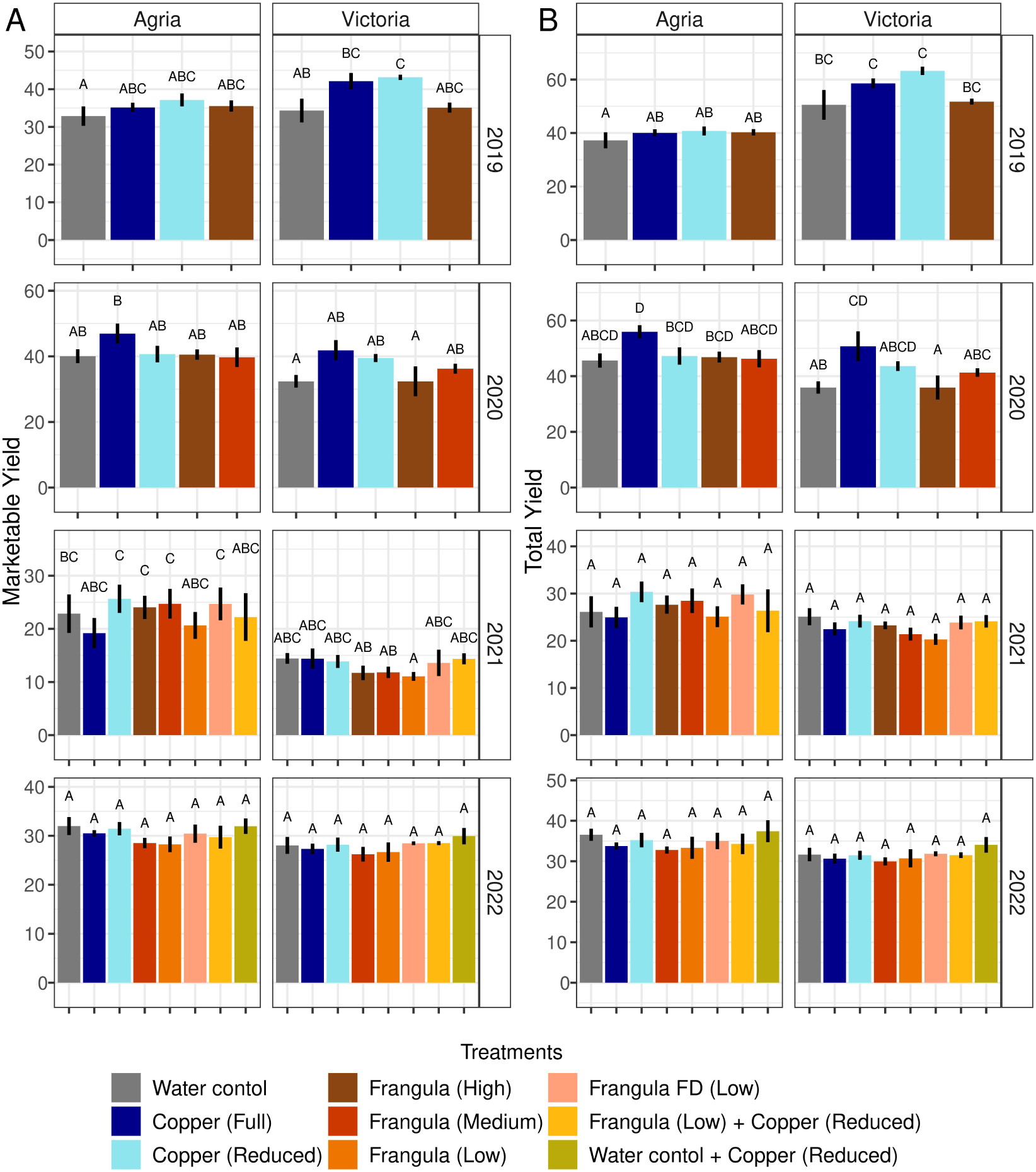
Yields from potato late blight field experiments between 2019 and 2022, including A) marketable yield (tons per ha) of tubers in two varieties (in size classes: 42-55 mm and 55-70 mm) and B) total yield (tons per ha) of tubers in two varieties of all size classes. Treatments are shown by color and described in Table 1. Bars indicate the standard error of the mean.

**Table 2:** ANOVA table of the linear mixed-effects models of experimental treatment, variety and their interaction on disease pressure and marketable and total yield from field trials between 2019 and 2022.

| Response variable | Year | Factor | Df (df1, df2) | F-value | p-value |
| --- | --- | --- | --- | --- | --- |
| AUDPC<sup>1</sup> | 2019 | Treatment | 3, 35 | 140.82 | < 0.001 |
|  |  | Variety | 1, 35 | 7.33 | 0.010 |
|  |  | Treatment x Variety | 3, 35 | 1.47 | 0.239 |
|  | 2020 | Treatment | 4, 45 | 63.43 | < 0.001 |
|  |  | Variety | 1, 45 | 0.76 | 0.389 |
|  |  | Treatment x Variety | 4, 45 | 0.69 | 0.605 |
|  | 2021 | Treatment | 7, 60 | 40.44 | < 0.001 |
|  |  | Variety | 1, 60 | 3.46 | 0.068 |
|  |  | Treatment x Variety | 7, 60 | 0.34 | 0.931 |
|  | 2022 | Treatment | 7, 60 | 28.68 | < 0.001 |
|  |  | Variety | 1, 60 | 7.08 | 0.010 |
|  |  | Treatment x Variety | 7, 60 | 0.83 | 0.565 |
| Marketable yield | 2019 | Treatment | 3, 35 | 4.99 | 0.006 |
|  |  | Variety | 1, 35 | 5.58 | 0.024 |
|  |  | Treatment x Variety | 3, 35 | 1.41 | 0.257 |
|  | 2020 | Treatment | 4, 45 | 3.65 | 0.012 |
|  |  | Variety | 1, 45 | 11.77 | 0.002 |
|  |  | Treatment x Variety | 4, 45 | 1.17 | 0.333 |
|  | 2021 | Treatment | 7, 60 | 0.53 | 0.81 |
|  |  | Variety | 1, 60 | 64.83 | < 0.001 |
|  |  | Treatment x Variety | 7, 60 | 0.83 | 0.568 |
|  | 2022 | Treatment | 7, 60 | 1.44 | 0.206 |
|  |  | Variety | 1, 60 | 10.78 | 0.002 |
|  |  | Treatment x Variety | 7, 60 | 0.24 | 0.975 |
| Total yield | 2019 | Treatment | 3, 35 | 3.62 | 0.023* |
|  |  | Variety | 1, 35 | 69.89 | < 0.001* |
|  |  | Treatment x Variety | 3, 35 | 1.13 | 0.351 |
|  | 2020 | Treatment | 4, 45 | 6.46 | < 0.001* |
|  |  | Variety | 1, 45 | 18.83 | < 0.001* |
|  |  | Treatment x Variety | 4, 45 | 1.25 | 0.303 |
|  | 2021 | Treatment | 7, 60 | 0.98 | 0.452 |
|  |  | Variety | 1, 60 | 12.45 | < 0.001* |
|  |  | Treatment x Variety | 7, 60 | 0.62 | 0.739 |
|  | 2022 | Treatment | 7, 60 | 1.31 | 0.263 |
|  |  | Variety | 1, 60 | 15.40 | 0.001 |
|  |  | Treatment x Variety | 7, 60 | 0.11 | 0.998 |

<sup>1</sup>Area under the disease progress curve
\*Significant after correction for multiple comparisons

At the time of harvest, tubers infected with late blight were found only in 2021 (0 to 4 infected tubers per treatment and variety were found in the 240-250 tubers analyzed per treatment and variety). There was no difference in the number of infected tubers across the treatments (*χ*^2^ = 10.88, df = 7, p-value = 0.144) or varieties (*χ*^2^ = 2.15, df = 1, p-value = 0.142). In all years, no tuber blight was found in the 3941 tubers examined one month after harvest from all treatments.

### 3.0.2 *Frangula alnus* extraction duration experiment on detached leaves

The detached Bintje leaves that had been treated with *F. alnus* were significantly less infected with *P. infestans* compared to the negative water controls (*χ*^2^ = 38.08, df = 5, p-value < 0.001). There was no difference between *F. alnus* treatments when the extraction time was 30 min or 120 min based on pairwise comparisons (Figure 4). Between the *F. alnus* treatments within the experiment, neither the treatment timing (one day, two days, and both one and two days prior to inoculation) (*χ*^2^ = 2.62, df = 2, p-value < 0.270) nor the extraction time (*χ*^2^ = 0.312, df = 1, p-value < 0.576) affected the infection rate. There was also no interaction between treatment timing and the *F. alnus* extraction time (*χ*^2^ = 2.44, df = 2, p-value < 0.295). Furthermore, the UHPLC profile of the *F. alnus* treatments did not show major differences in extracted compounds between the different extraction times, although quantities of compounds extracted differed (see section below on the chemical analysis of *F. alnus* for more details). Because no difference was found in treatment efficacy based on extraction time, the extraction time for *F. alnus* treatments in the field experiments was reduced to 30 minutes starting in 2021.

**Fig. 4:**
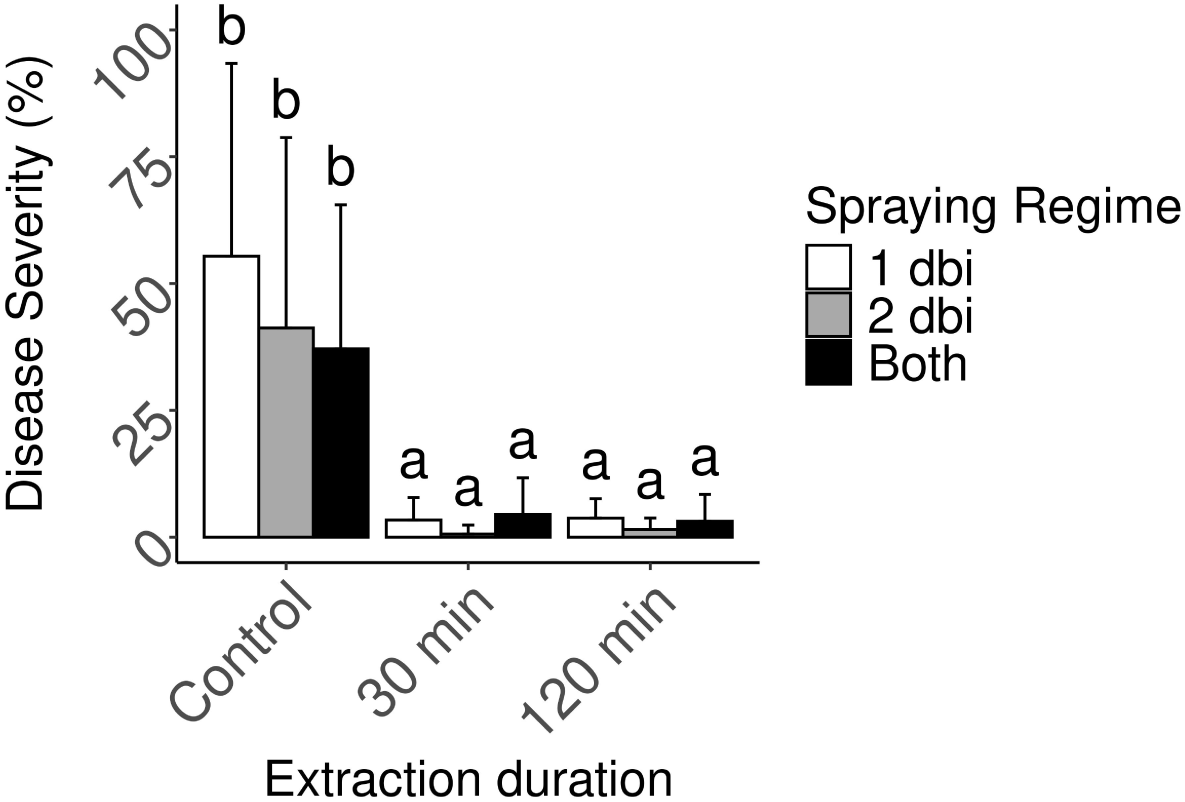
Disease severity on detached leaves when sprayed with *Frangula alnus* that had been extracted in an aqueous solution for 30 or 120 minutes compared to a negative control treatment of tap water. The treatments included spraying regimes consisting of one day before, two days before, and both one and two days before inoculation (dbi) of *Phytophthora infestans*. Bars indicate the standard error of the mean.

### 3.0.3 *F. alnus* dosage experiment on detached leaves

ased on detached leaf assays, the Kruskal- Wallis test revealed a difference among *F. alnus*extracts and resulting disease severity (*χ*^2^ = 33.98, df = 5, p-value < 0.001). Dosages less than 2.5 kg/ha (Figure 5) were significantly less effective than the dosages that were equal to or greater than 2.5 kg/ha. There were no significant differences in disease severity among the highest three dosages (2.5, 25 and 250 kg/ha). The disease severity found in the treatments of the lowest two dosages (0.025 and 0.25 kg/ha) did not differ from the water- control. The IC50, indicating the dosage at which *F. alnus* inhibits half of the *P. infestans* infection that it is maximally capable of, was found to be 1.253 kg/ha. At 2.5 kg/ha, 69.4% of the maximum inhibitory potential of *P. infestans* was reached based on this dosage experiment.

**Fig. 5:**
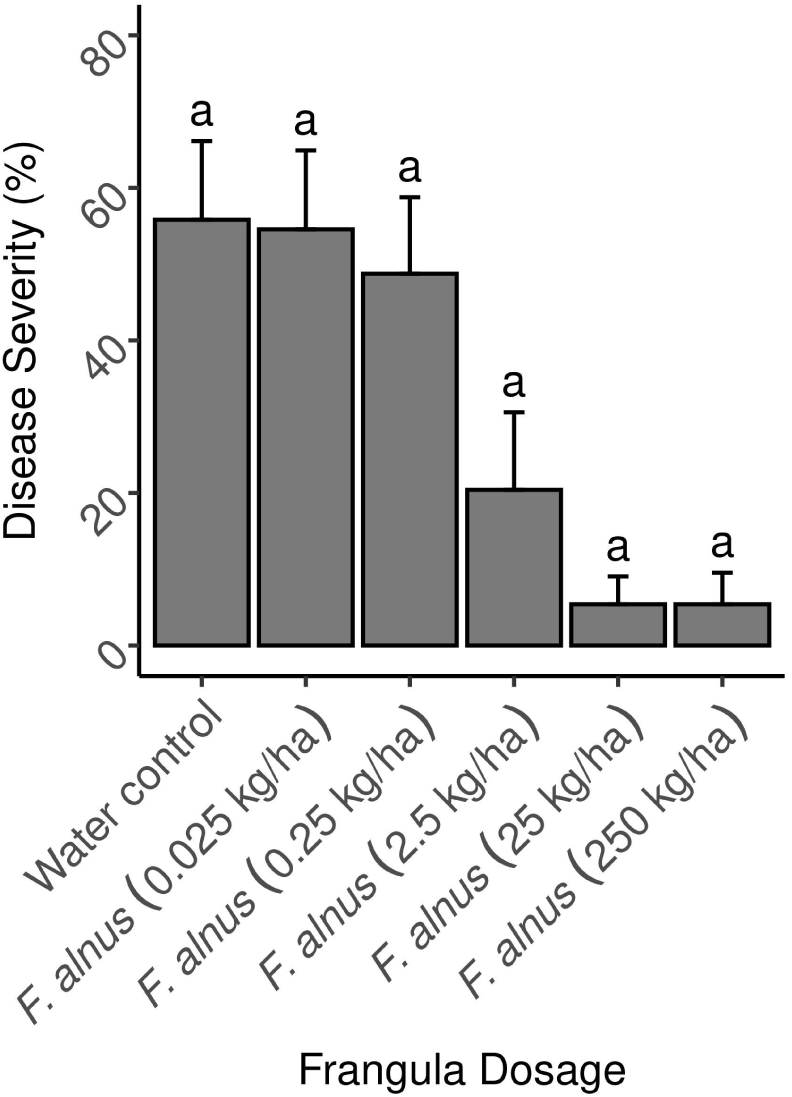
Disease severity of detached leaves when sprayed with different dosages of *Frangula alnus* compared to a negative control treatment of tap water shown with the standard error of the mean. The 25 kg/ha dosage corresponds to the medium dosage from the 2020-2022 field experiments while 2.5 kg/ha is the same dosage applied to the low *F. alnus* treatment in the field (Table 1).

### 3.0.4 *In vitro* mycelial growth experiment

The mycelia of *P. infestans* showed significantly different levels of growth inhibition depending on the treatment (*χ*^2^ = 88.35, df = 10, p-value < 0.001). There were no differences in mycelial growth on plates that received the autoclaved deionized water control, and the filtered *F. alnus* low treatment, and both filtered and non-filtered *F. alnus* freeze-dried (FD) treatments (Figure 6). For the high, medium, and low *F. alnus* treatments that had been prepared by aqueous extraction, there was significantly greater mycelial inhibition compared to their respective filtered treatments of the same dosages. The non-filtered medium and high *F. alnus* treatments that resulted in the growth of the *Erwinia* spp. inhibited mycelial growth more than the filtered treatments of the same dosages (Figure 7). For filtered treatments prepared with an aqueous extraction, the low *F. alnus* dosage showed decreased efficacy compared to the medium and high *F. alnus* treatments. The full copper treatment showed an increase in mycelial inhibition compared to the reduced copper treatment but no difference from the medium and high non-filtered *F. alnus* treatment where bacterial colonization of the *Erwinia* spp. was observed.

**Fig. 6:**
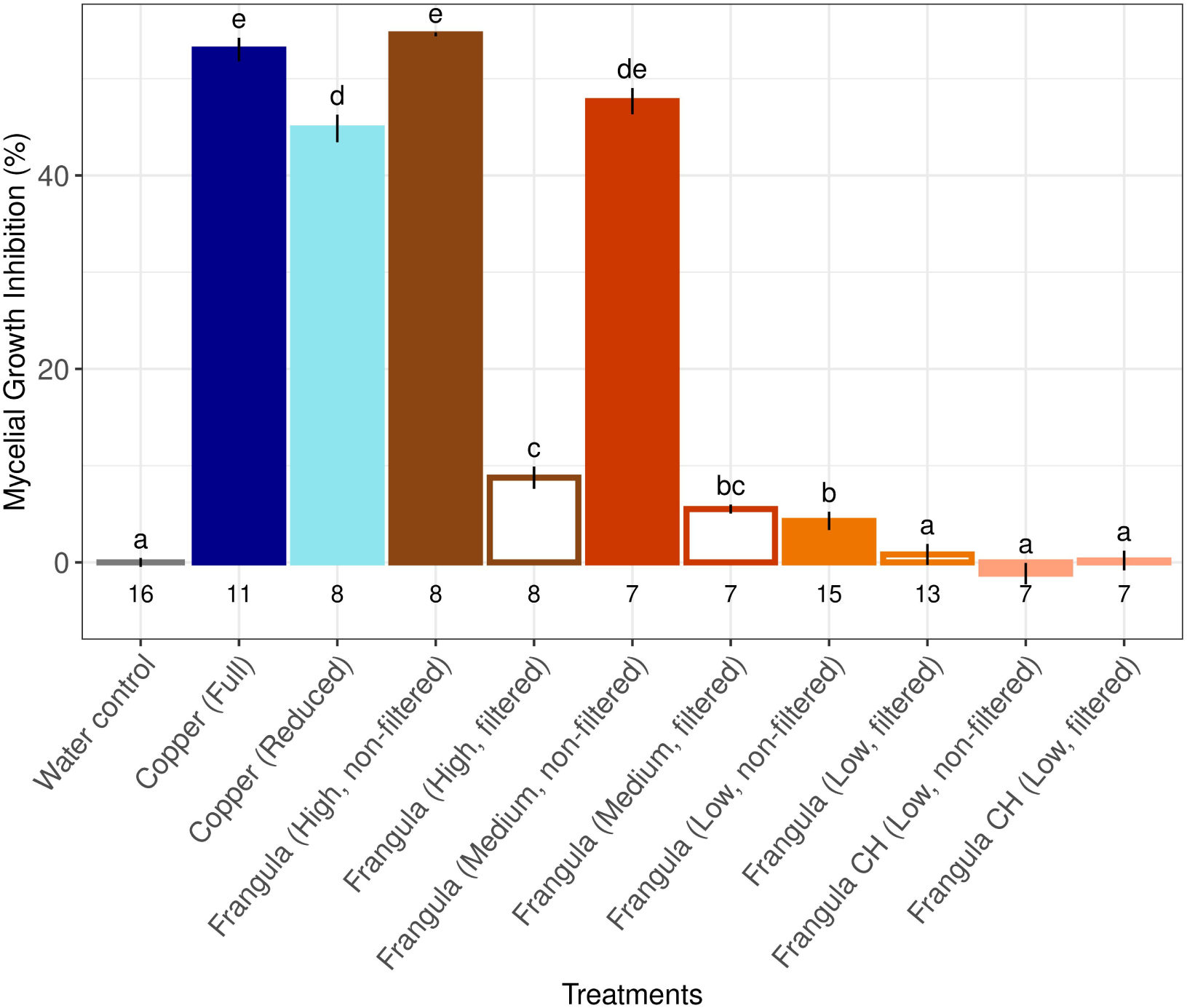
Mycelial growth inhibition in vitro experiments on rye agar. Measurements of the mycelial area of all treatments were compared to the negative control that had received autoclaved water. The reduction of mycelial area shows the growth inhibition due to the direct efficacy of the treatment on *Phytophthora infestans*. Treatment descriptions are found in Table (1). The numbers below the x-axis represent the number of plates included in each treatment. Bars indicate the standard error of the mean.

**Fig. 7:**
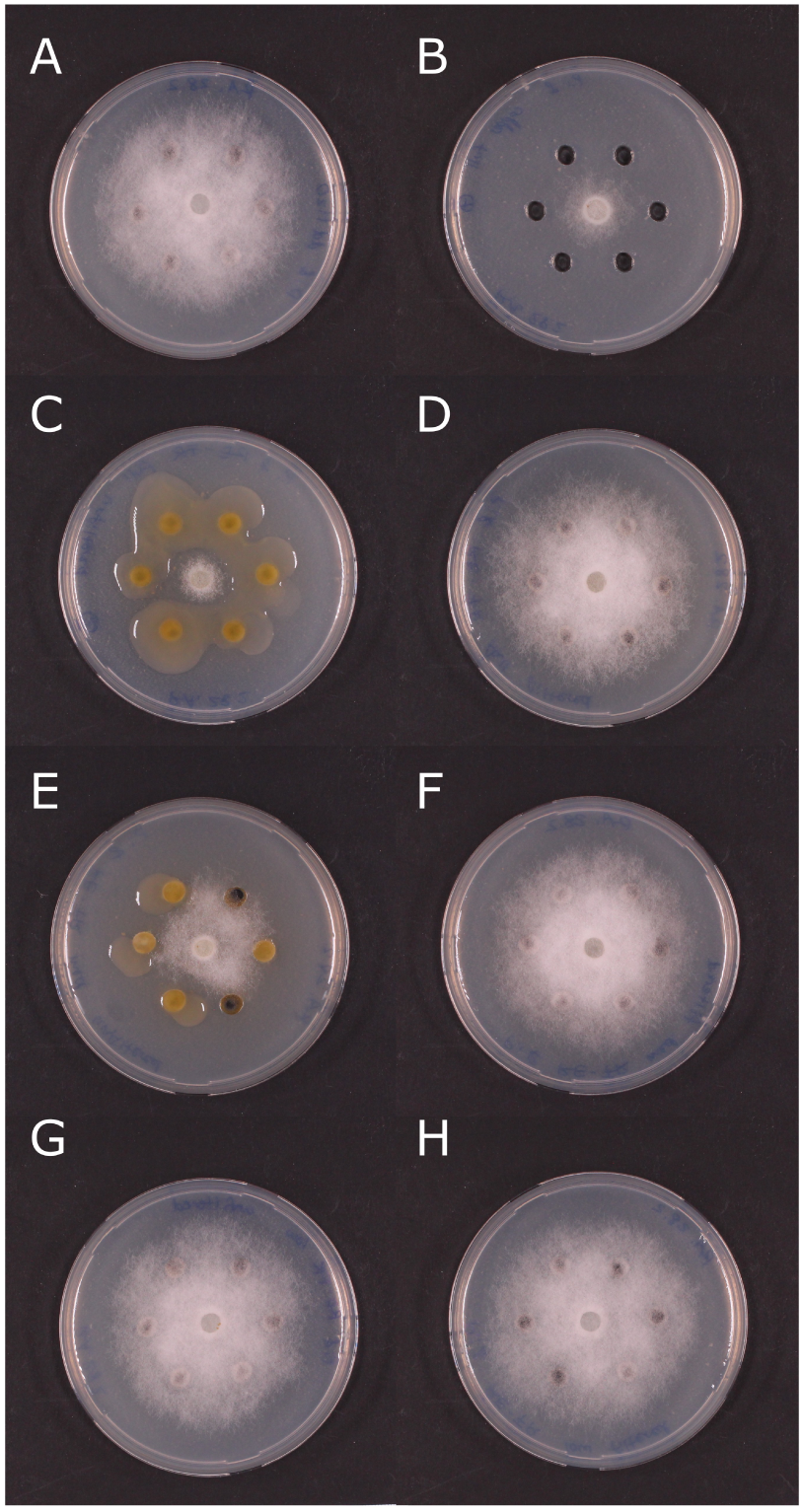
Pictures from in vitro assays to assess treatment inhibition of *Phytophthora infestans* mycelial growth. Treatments shown include a) negative water control b) copper (full) dosage c) *Fragnula alnus* high dosage, non-filtered d) *F. alnus* high dosage, filtered e) *F. alnus* medium dosage, non-filtered f) *F. alnus* medium dosage, filtered g) *F. alnus* low dosage, non-filtered h) *F. aluns* low dosage, filtered. The bacterium associated with the *F. alnus* bark that colonized the plates in treatments shown in panels c) and e) was identified to be an *Erwinia* spp.

### 3.0.5 Detached leaf assay with filtered *F. alnus*

A detached leaf assay (assay A) was run to test the difference between the filtered and non-filtered *F. alnus* treatments that had been prepared with an aqueous extraction method. The assay was then run a second time (Assay B) to determine the consistency of the results. There were differences among treatments in both assay A (*χ*^2^ = 29.133, df = 3, p-value = 0.001) and assay B (*χ*^2^ = 35.648, df = 3, p-value = 0.001). The autoclaved deionized water control showed significantly higher rates of disease severity in both assays (Figure 8), while the copper treatment yielded a significantly lower disease severity than the *F. alnus* treatments in assay A. In assay B, copper and *F. alnus* treatments showed either low or no *P. infestans* infections, and the treatments were not different from each other.

**Fig. 8:**
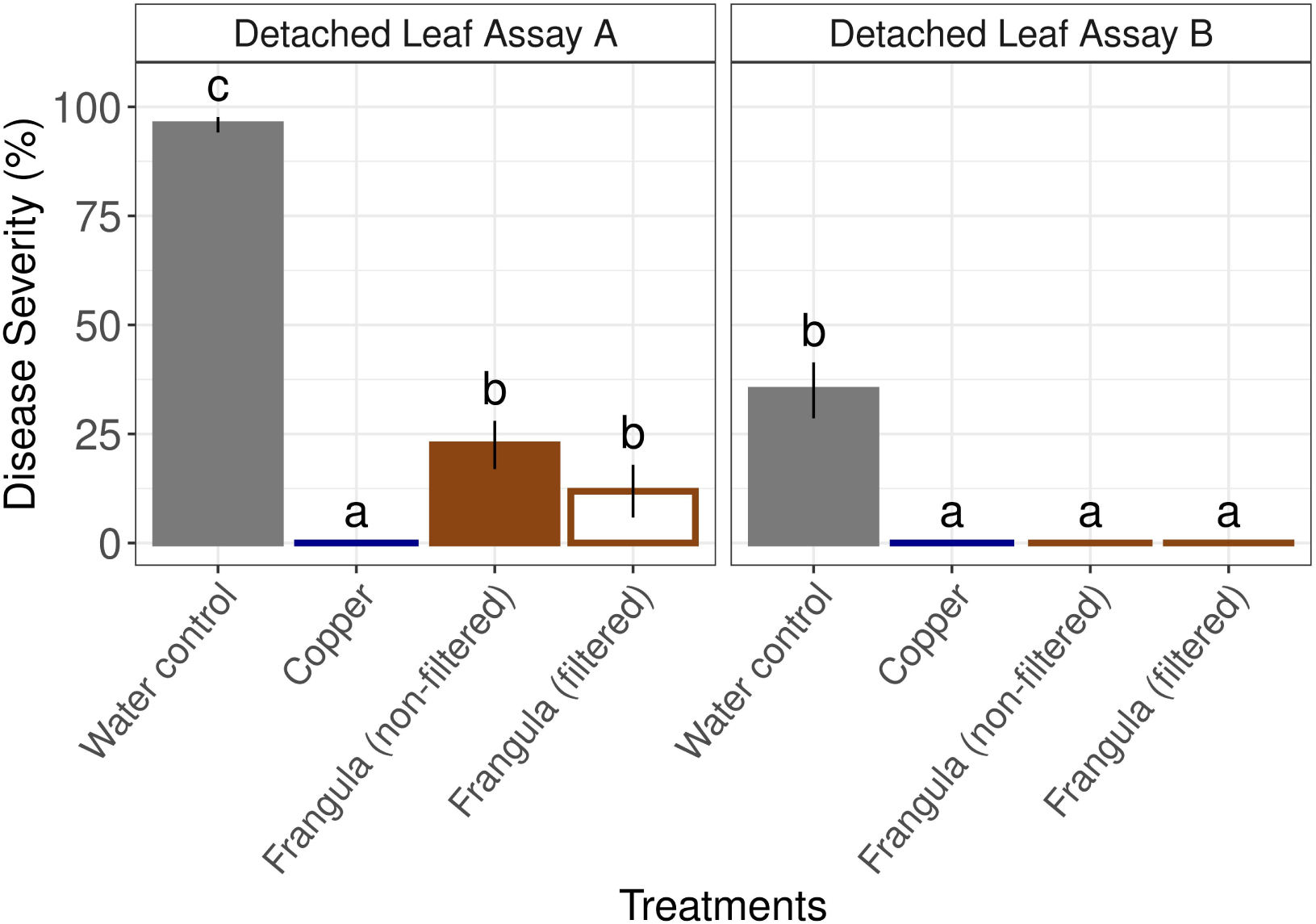
Two assays (Assay A and Assay B) were run on detached leaves in which *Frangula alnus* that had been filtered was compared to *F. alnus* that had not been filtered. The high dosage of *F. alnus* was used for this experiment. Copper was included as a positive control. Bars indicate the standard error of the mean.

### 3.0.6 Chemical analysis of *Frangula alnus* extracts

The Ultra High Performance Liquid Chromatography—charged aerosol detector (UHPLC-CAD) profile of the different dosages of the aqueous suspension of F. alnus (medium and low dosage) showed a similar chemical composition, differing only on the concentration of the extracted compounds. However, the FD-low dosage showed that the filtering and freeze-drying step concentrates the extract on polar compounds, thus reducing the relative quantity of semi-polar and apolar compounds (i.e. anthraquinones). Indeed, the quantification of the main constitutive anthraquinones differed among *F. alnus* dosage amounts (Figure 9, Frangulin A: *F*_(2,8)_ = 299.190, p < 0.001, Frangulin B: *F*_(2,8)_ = 205.947, p < 0.001, Emodin: *F*_(2,8)_ = 20.928, p < 0.001). As expected, the medium dosage consistently yielded more anthraquinones than the low dosage. The medium dosage included 10X more ground *F. alnus* bark than the low dosage, but it proportionally yielded more Frangulin A and Frangulin B (17-18X more) than the low dosage, and proportionally less Emodin (only 5X more). The low dosages with the lyophilization step (FD-Low) also yielded comparatively lower compounds than both aqueous extractions (2.5X less than the low dosage). Although the duration of the aqueous extraction did not change the efficacy of the *F. alnus* treatments (Figure 4), it seems as though the 30 min extraction yielded more Frangulin A and Frangulin B than the 120 min extraction but similar levels of Emodin (Figure A4). Based on two measurements of the *F. alnus* preparation following both extraction periods, the 30 min extraction yielded on average 4 times more Frangulin A (100.4 *µ*g/mL versus 23.5 *µ*g/mL) and almost 3 times more Frangulin B (52.7 *µ*g/mL versus 18.2*µ*g/mL) than the 120 min extraction.

**Fig. 9:**
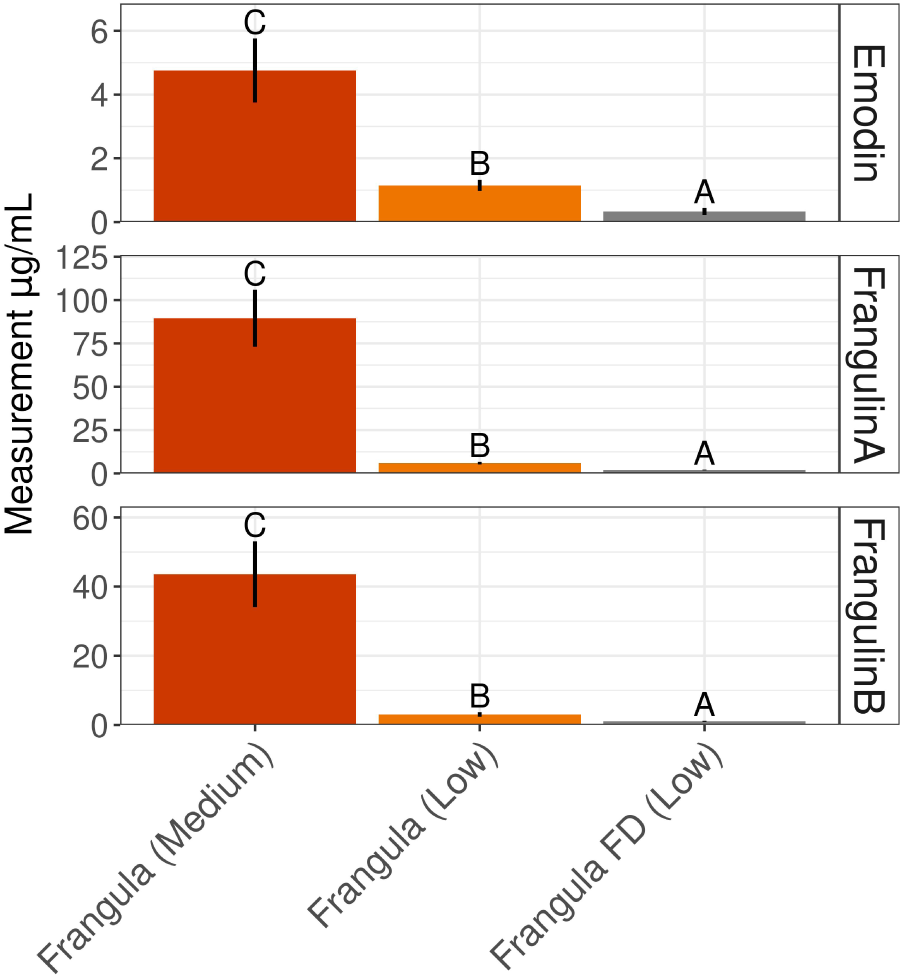
Chromatographic measurements of compounds extracted from *Frangula* preparations that were associated with potato late blight reductions. The medium dosage corresponds to 25 g/ha and the low dosage to 2.5 g/ha from the ground bark. The FD extraction included a lyophilization step while the other extractions were stirred at room temperature in tap water for 30 min. Bars indicate the standard error of the mean.

## 4 Discussion

Phytophthora infestans is difficult to control due to its ability to evolve and evade host resistance (Leesutthiphonchai et al, 2018). In countries that still register copper as a plant protection product, it effectively controls PLB on organic farms due to its multi-site mode of action (La Torre et al, 2018). However, because its residues persist and can detrimentally affect soil organisms, countries that still allow its use seek avenues to reduce it (Silva et al, 2022; Möhring et al, 2020). In 12 European countries, 56% of producers already use half of the allowable amount of copper (Tamm et al, 2022), showing that the reduction is feasible and could potentially be more broadly adopted.

Our research contributes additional evidence that shows using only half of the allowable copper (2 kg/ha/year) compared to the full allowable amount in Switzerland (4 kg/ha/year) can result in similar levels of disease control and potato yield. Previous work also showed that reduced copper did not tend to affect yield, which was rather driven by year, site, variety, and disease pressure (Wiik, 2014; Gonźalez-Jiménez et al, 2023). In our experiments, variety influenced marketable and total yield more than treatment. In 2021, the increased disease pressure reduced the yield for both varieties, especially for Victoria. The Agria plots in our study had a lower yield reduction (35%) than the average yield reduction estimated across organic production (57%) in 2021 compared to yields in 2019, 2020 and 2022 (Bio Suisse, 2022). On the other hand, the yields in our Victoria plots were reduced by 63%. These varietal differences in yield reduction underscore the role that potato variety selection has in combating PLB’s damage; the yield differences between varieties were much greater than any treatment differences in our study.

Treatment had a limited effect on yield and no effect on tuber infections, which may have been due to the relatively late start of the epidemic or low disease pressure in three of the four years (Wiik, 2014). The plot size of the experiment was smaller than recommended to measure yield differences (EPPO, 2021), which may also play a role in the outcome. The disease severity as measured by AUDPC, on the other hand, fluctuated to a much greater degree among treatments. The years with low to moderate disease pressure in the field experiments, including 2019, 2020 and 2022, the *F. alnus* treated plots had lower disease severity compared to the water control. The reduction from the *F. alnus* medium and high dosages in the 2020 field experiment confirmed results from former field studies ran in 2010-2012 (Forrer et al, 2017). In these experiments, there was no difference in PLB disease severity between a 4% *F. alnus* solution and a reduced copper treatment (0.3 kg/ha per treatment applied 8 times), and the *F. alnus* treatment reduced disease severity by 37% based on the AUDPC.

During the last two years of our experiments, we no longer included the *F. alnus* high dosage treatment due to the lack of economic feasibility. Instead, a low dose, consisting of 2.5 kg/ha was included, which would be in the economic range of copper and fungicide treatments (430-580 CHF per season/ha). The low dosage treatment can inhibit 69.4% of the pathogen compared to the maximal inhibitory concentration of *F. alnus* based on the laboratory-based dosage experiment. During a high disease pressure year such as 2021, however, the inhibition is not effective at reducing disease severity in the field based on the AUDPC. Because the *F. alnus* extraction has not undergone any subsequent formulation, its leaching may also have contributed to its inability to inhibit *P. infestans*. Nonetheless, it reduced disease severity based on the AUDPC in 2022 along with all *F. alnus* dosages, although it was a year with minimal PLB pressure.

In 2021, only copper-containing treatments showed a significant reduction in PLB, including one treatment (treatment 8) that switched from low dosage *F. alnus* to copper treatments on the fifth spraying application, resulting in overall less copper use. This sequential treatment, during which a *F. alnus* solution was first sprayed until disease pressure was elevated, followed by a switch to a low dose copper treatment mid-way through the season, , resulted in a 70% reduction in copper application compared to the full dosage (i.e. the maximum application allowed in Switzerland per year with 10 applications). Therefore, delayed application of reduced dosages (treatments 8 and 9) could be an effective means to reach targets of reduced copper applications. Further work is needed to demonstrate whether the *F. alnus* gives an added benefit or copper’s delayed application is sufficiently protective without any early *F. alnus* treatments. However, *F. alnus* clearly reduces *P. infestans* infection as demonstrated through the field studies in years of low to moderate PLB severity and reinforced by our laboratory experiments. Previous work has focused on mixed treatments that are sprayed on the same date (Liljeroth et al, 2010; Stridh et al, 2022), and there is less information about sequentially used treatments.

The laboratory studies provided a controlled environment to gain further information about the mode of action and effect of different spraying regimes and application dosages. The in vitro work was performed to understand if *F. alnus* directly inhibits P. infestans mycelial growth. When the non-filtered *F. alnus* aqueous extractions were plated, the *Erwinia* spp. associated with the *F. alnus* bark established on the plate and directly inhibited or competed with *P. infestans*. The bacterium did not grow from the *F. alnus* extracted through lyophilization (treatment 7, low dose, FD), the filtered extracts nor on the copper and water control plates. The effect of the bacterium originating from the *F. alnus* bark on *P. infestans* in the in vitro experiment also suggests that botanical applications may act not only through the chemical compounds from the plant, but depending on the extraction method, potentially also through their microbiota. The microbiota of plants can provide protection against pathogens (Vogel et al, 2021), and perhaps the application of botanical plant protection products could also transfer protective microbial constituents through targeted mycobiome transplants (Chock et al, 2021). However, it is unclear to what extent the microbiota of *F. alnus* treatment actually establishes on the potato plant, and further work would need to directly assess the colonization ability of a treatment’s microbiota. Additionally, it is also unclear how widespread *Erwinia* spp. co-occurs in *F. alnus* bark. Previous work, in which F. alnus extracts were plated mentioned no bacterial colonization (Forrer et al, 2017). Nonetheless, the possible transmission of microbiota could potentially influence a botanical treatment’s effect and should be considered in future studies.

To disentangle the inhibition from the bacterium and *F. alnus*’s chemical compounds, the extracts from all treatments were added to the in vitro assays either after filtration to eliminate the bacterium or directly without filtration. These in vitro results suggest that the chemical compounds themselves provided minimal direct inhibition to *P. infestans* mycelial growth compared to the bacterium associated with the bark, which appears to provide most of the in vitro inhibition. These findings corroborate previous results that showed *F. alnus* only slightly inhibited mycelial growth (Forrer et al, 2017). However, it was found that F. alnus can inhibit sporangial germination of P. infestans (Forrer et al, 2017). Indeed, several anthraquinones and their analogues have been shown to possess antifungal activity against several phytopathogens (Kim et al, 2004; Barilli et al, 2022; Choi et al, 2004), and the presence of these compounds in the F. alnus extracts might account for decreased *P. infestans* through direct mechanism such as decreased sporangial germination and slightly decreased mycelial growth.

Unlike in vitro tests, both filtered and unfiltered *F. alnus* treatments, resulted in significantly reduced *P. infestans* disease severity on detached leaves. The filtered *F. alnus*, which had a much lower direct effect on *P. infestans* in vitro, showed as much efficacy as the non-filtered extraction in detached leaf assays, pointing to the possible involvement of induced resistance (Forrer et al, 2017). Anthraquinones have also been shown to induce resistance in plants (Konstantinidou-Doltsinis and Schmit, 1998), and other chemical families present in the F. alnus extracts, such as polyphenols, have been implicated in defense induction (Gillmeister et al, 2019). In *Vitis vinifera*, the application of *F. alnus* bark extract induces the production of the phytoalexins piceid, resveratrol and pterostilbene, and protects against infection of the oomycete *Plasmopara viticola* (Godard et al, 2009). Plant elicitation can complement direct antifungal activity if deployed in the right situations to avoid reduced yields (Heil et al, 2000), and therefore, genotypic and environmental variables should be evaluated before applying elicitors since these factors can influence their success (Walters et al, 2013). For example, previous research with F. alnus showed a varietal difference in yield (Forrer et al, 2017), which may be due to the way in which a potato genotype responds to an elicitor (Bruce, 2014) and may also account for the varietal differences we see in yield in our field experiments.

Current obstacles to adoption include cost of the material, effort required for the preparations, and stability of the extraction. To further develop an effective and potentially cost-effective treatment, more information about the compounds involved could better elucidate the mode of action. Frangulin A, Frangulin B and Emodin were found in the *F. alnus* extract, but it is not clear which chemicals are responsible for the direct or indirect inhibition. Singularly testing each compound would help decipher how they individually and cumulatively control PLB. These steps could help determine whether a stable formulation can be achieved to obtain a cost-effective treatment. Therefore, many steps are still required (i.e. an understanding of kinetic degradation of the active compounds, stability in solution and on foliage, optimal formulations) to achieve a viable PLB spraying regime that incorporates *F. alnus* or its components.

Our work suggests that copper application amount could be substantially reduced based on the low dose copper treatment’s results that showed no yield differences from the full dose treatment. Potentially a low dose copper treatment could be supplemented with alternative products, such as *F. alnus*, but it is not yet clear if the botanical gives added value. Further testing of sequential deployments of different treatments (i.e. switch from an alternative treatment, such as a botanical or another plant elicitor to copper) can also help contribute to overall copper reduction. The success of such strategies would be dependent on the response of the variety to plant elicitation, and the timing of the epidemic may also very much impact their success. Reductions in copper application combined with the deployment of resistant varieties (Kessel et al, 2018) can help achieve goals of reduced pesticide targets (Silva et al, 2022; Purnhagen et al, 2021). Due to importance of potato production in Europe a combination of strategies are necessary to combat potato late blight.

## Supplementary information

Detailed weather and agronomic information for all four years of the field experiment as well as additional information about sample and data analyses can be found in the supplement.

## Author contributions

The study was conceived and designed by Tomke Musa, Karen Sullam, and Andreas Kägi. Material preparation, data collection and analysis were performed by Tomke Musa, Karen Sullam, Haruna Gütlin, Andreas Kägi, Sylvain Schnee and Josep Massana-Codina. Tomke Musa and Karen Sullam jointly wrote the first draft of the manuscript. All authors commented on previous versions of the manuscript and read and approved the final manuscript.

## Acknowledgments

We thank Susanne Vogelgsang for the valuable support and constructive feedback. We also thank Damian Oswald, Nicole Togni, Lisa Besson, Arnaud Chotel, Anja Logo, Cecilia Panzetti, and Clara Chevalley for their assistance with laboratory and/or field experiments.

## Appendix A Supplementary Material

### A.1 Agronomic and meteorological conditions

In order to clarify the conditions and set-up used in the field experiments, we provide the following supplementary information, including information on the plot set-up (Figure A1), weather during the field experiment (Figure A2), and agronomic conditions and treatment applications (Tables A1 and A2).

**Fig. A1:**
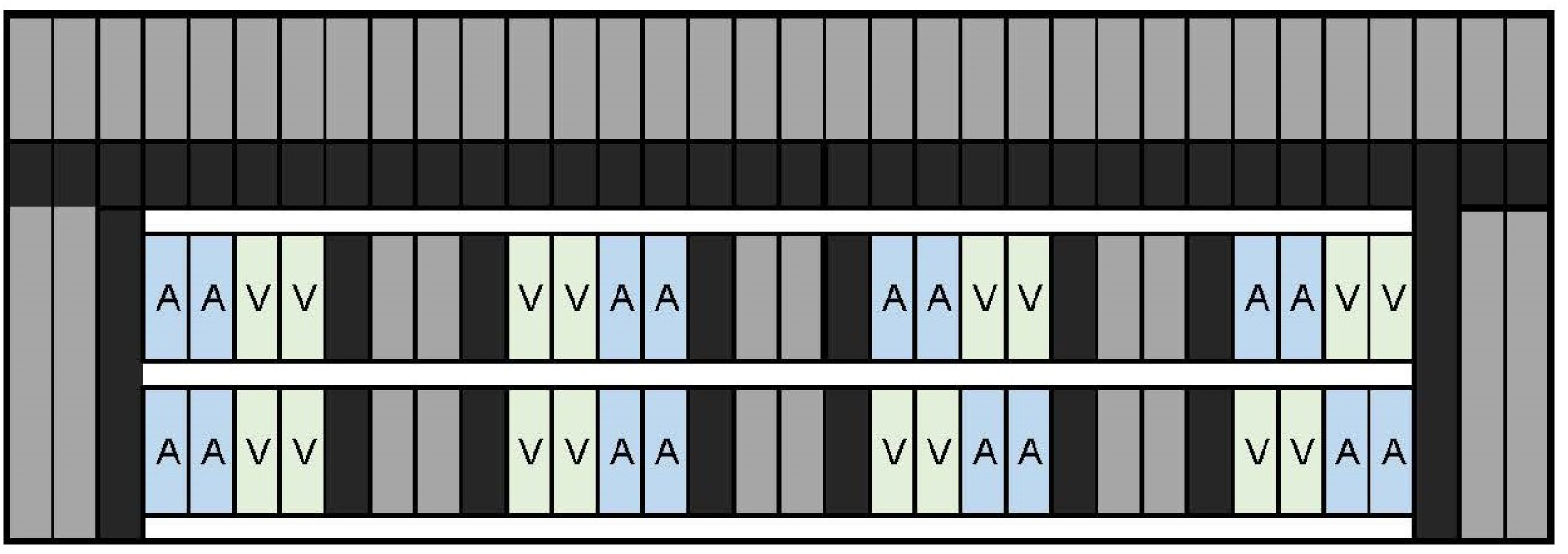
Schematic diagram of the potato variety split-plot design used in the field experiment. Each sub-plot included 2 rows of one variety (Agria or Victoria) next to the other variety. These plots were sandwiched between a low susceptible variety (in black) and the high susceptible variety in gray. Each box represents a different row.

**Fig. A2:**
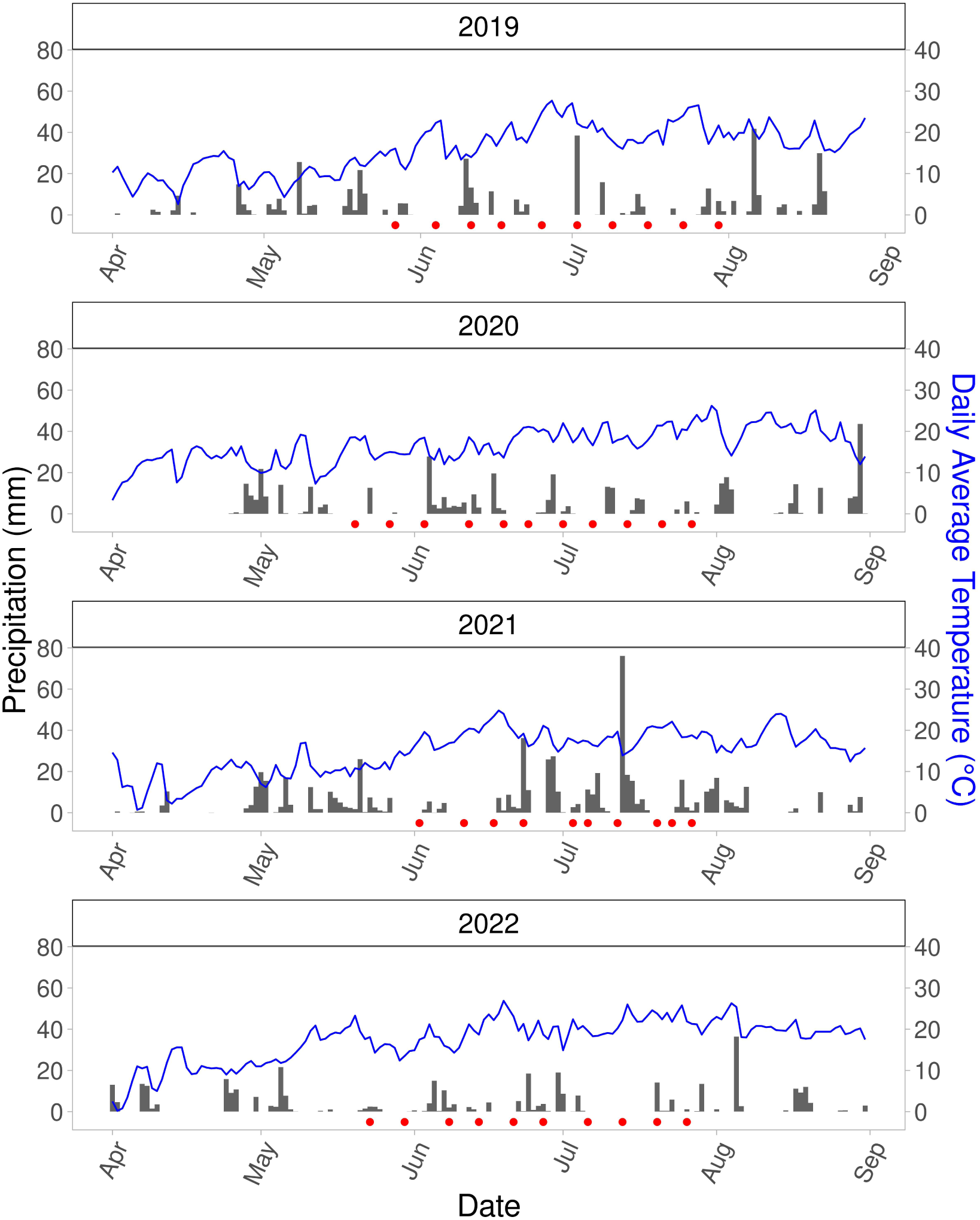
Graphs of meteorological conditions during the field experiments between 2019 to 2022, showing the daily average temperature (°C) with blue lines (y-axis on the right side) and daily sum of precipitation (mm) with gray bar (y-axis on the left side). The dates during which treatments were applied are shown by the red point along the x-axis. Generally, the treatments were applied in the mornings unless rain postponed the treatment. The weather measurements were taken at the MeteoSwiss station Zurich/Affoltern.

### A.2 Identification of bacterial isolated from *F. alnus* aqueous extracts

The bacterial isolate was directly amplified with the primers 27F and 1492R (Heuer et al, 1997). Reactions were run with Taq-&Go^TM^ Mastermix (MP Biomedicals, Heidelberg, Germany) in a reaction volume of 20 *µ*L and primer concentration of 0.5 *µ*M. Reaction conditions included an initial denaturation at 94 °C for 5 min followed by 32 cycles of 94 °C 30 sec, 55 °C for 1 min and 72 °C for 1.5 min and a final elongation step of 72°C for 5 min. Sanger sequencing was performed (Microsynth, Balgach, Switzerland) with both 27F and 1492R, and a contig was constructed using Geneious Prime^®^ 2024.0.3. The sequence will be deposited in the NCBI nucleotide database (Accession #, to be provided).

### A.3 Additional details on *in vitro* and detached leaf experiments

For detached leaf experiments, terminal leaves, consisting of three leaflets were used (See Figure A3). The chromatograms from the detached leaf experiment using 30 min and 120 min extraction times of *F. alnus* are also displayed (See Figure A4).

**Table A1:** Agronomic information about field and environmental conditions during the four years of field experiments included in this study.

| **Agronomic Information** | **2019** | **2020** | **2021** | **2022** |
| --- | --- | --- | --- | --- |
| Field site | Rümlang, Plot 153 | Reckenholz, Plot 114A | Reckenholz Plot 119 | Reckenholz Plot 159 |
| Coordinates | 47.438796, 8.524827 | 47.430933, 8.520306 | 47.428847, 8.515277 | 47.433979, 8.511500 |
| Soil type | sandy loamy, very deep | clay, sandy loam | clay, clayey loam | clay, clayey loam |
| Pre-crop | buckwheat | canola | grass seed propagation | summer wheat, intercrop (green manure) |
| Soil management | 12 August 2018 cultivator, 18 April 2019 harrowing | 6 April 2020 harrowing | — | 18 April 2022 harrowing |
| Fertilization | compost, total per ha: 68 N, 152 P, 315 K, 0 Mg | total per ha: 112 N, 0 P, 360 K, 26 Mg | total per ha: 106 N, 87.4 P, 120 K, 36 Mg | compost, Patentkali, Super Urea, total per ha: 83 N, 222 P, 425 K, 145 Mg |
| Herbicide treatment | 7 May Bandur Boxer Sencor | 17 April Bandur Boxer Sencor SC; 28 May Fusilade Max | 30 April Stomp Aqua + Firebird Plus (some herbicide damage); 31 May Fusilade Max | 10 May Boxer; Sencor; 30 May Fusilade Max |
| Irrigation | — | — | — | 22 June, 24 June, 19 July |
| Control of Colorado beetle | 24 June (Biscaya) | 8 June (Biscaya) | 17 June (Audienz), 10 July (Biscaya) | 19 May (Audienz), 10 June (Audienz) |
| Planting | 18 April | 8 April | 20 April | 20 April |
| Emergence | 8 May | 1 May | 21 May | 10 May |
| BBCH15 | 20 May | 11 May | 26 May | 20 May |
| Defoliation of Bintje spreader rows | — | 6 July | 12 July | — |
| Defoliation of test varieties | 4 August | 27 July | 29 July | 2 August |
| Harvest | 29 August | 19 August | 18 August | 23 August |
| Date yield measurement | 16 September | 21 August | 29 August | 21 August |
| First late blight reported in Switzerland | 20 May | 11 May | 21 May | 10 May |
| Location of first announced PLB in Switzerland | Schlatt (TG), 30 km distance | Gürbrü (BE)/Kerzers (FR), 110 km distance | Dorf (ZH), 23 km distance | Flaach (ZH), 19 km distance, Weite (SG), 109 km distance |
| First late blight in field experiment | 11 June | 24 June | 6 July | 30 June |
| Overall weather conditions during field season | wet | wet | very wet | dry |
| Sum of precipitation (mm) during entire spraying period | 194.0 | 190.6 | 345.6 | 155.6 |
| PLB Pressure in Switzerland | low | medium | high | very low |

**Table A2:**
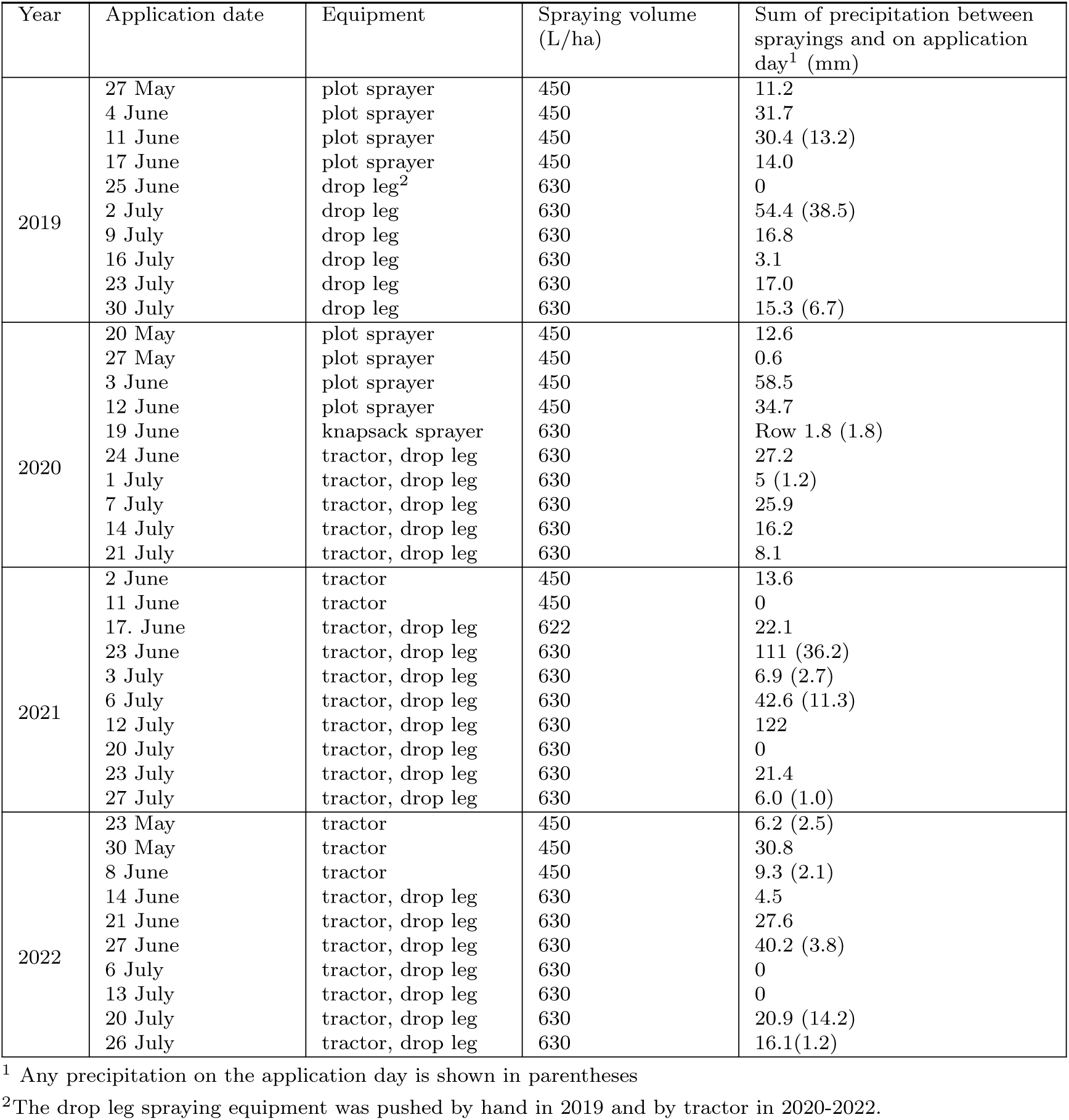
Dates of treatment application, equipment used for the application, volume of water used for the treatment mixtures (dry ingredient amounts shown in Table 1), and the sum of precipitation (mm) between spraying dates as well as on application day in parentheses.

**Fig. A3:**
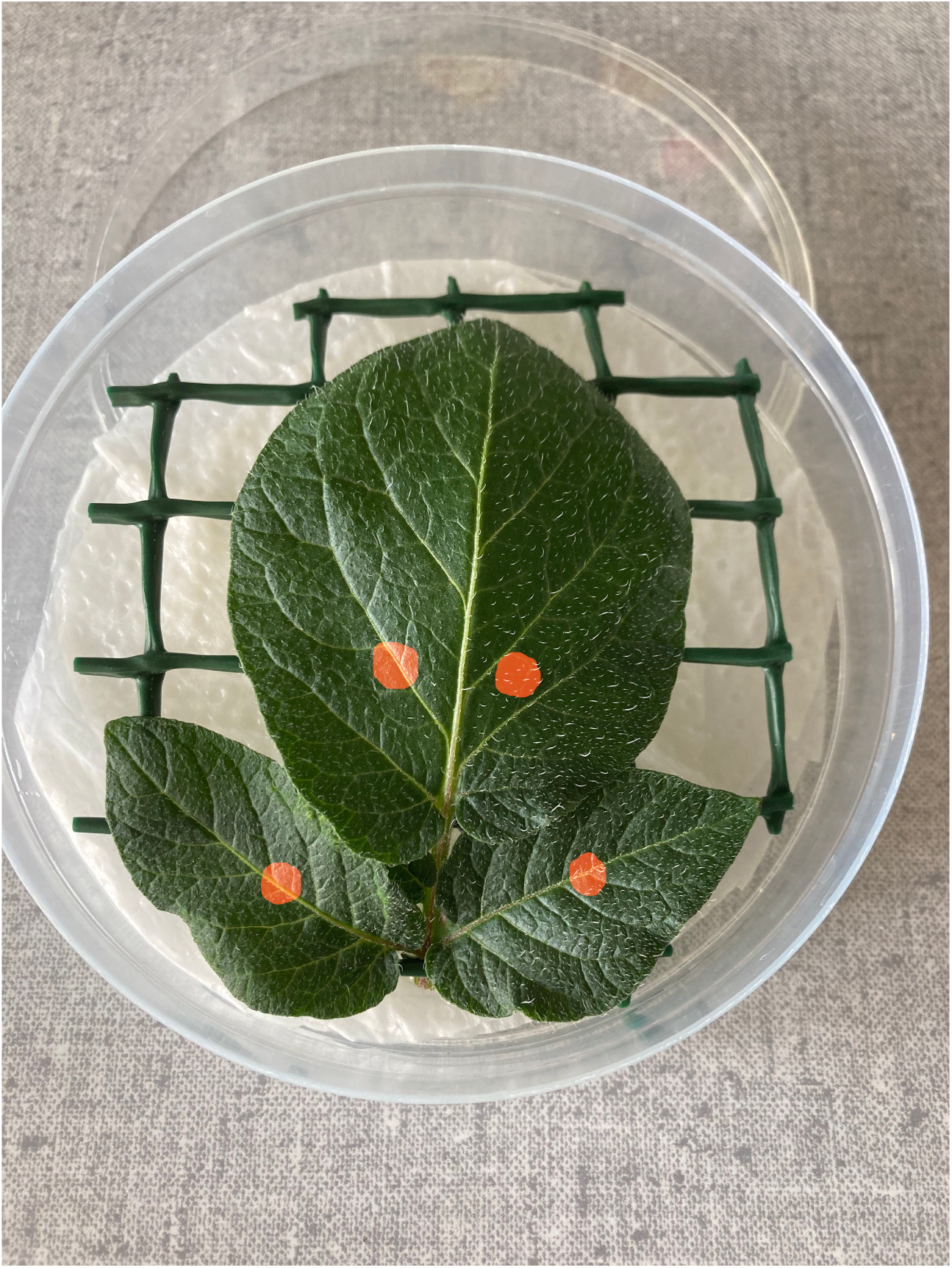
Each terminal leaflet was inoculated twice, and each of the two adjacent leaflets were inoculated once. The four locations on which 30 *µ*L of spore suspension were deposited for the *Phytophthora infestans* inoculations are indicated by the red markings in the picture. Leaves were placed on moistened paper towels and a wire rack in Petri dishes.

**Fig. A4:**
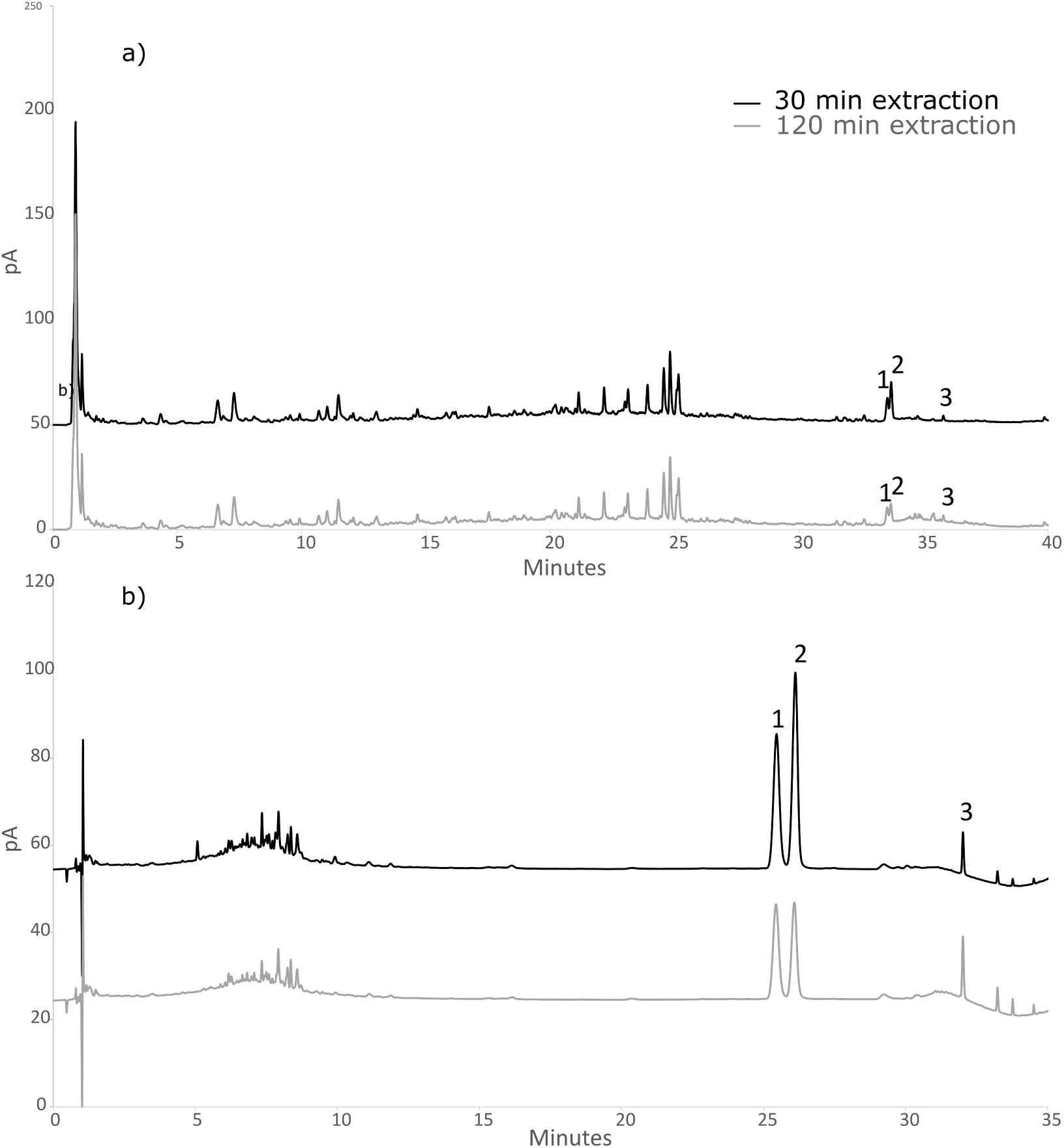
CAD (A) and UV at 435 nm (B) chromatograms of the *Frangula alnus* bark aqueous extract after 30 minutes and 120 minutes of stirring. The numbers associated with peaks in the chromatograms correspond to the following compounds: 1—Frangulin B, 2—Frangulin A, and 3—Emodin.

#### A.3.1 Image J Macro scripts

The following Image J macro was modified from a version previously published (Laflamme et al, 2016) to quantify the effect of *F. alnus* in the in vitro mycelial growth experiment. It automatically measures the area in a defined color threshold and saves a painted picture with the selected area for visualization. Adjusted measurements of the Petri dish and the mycelium were used to calculate the percentage of mycelial coverage.

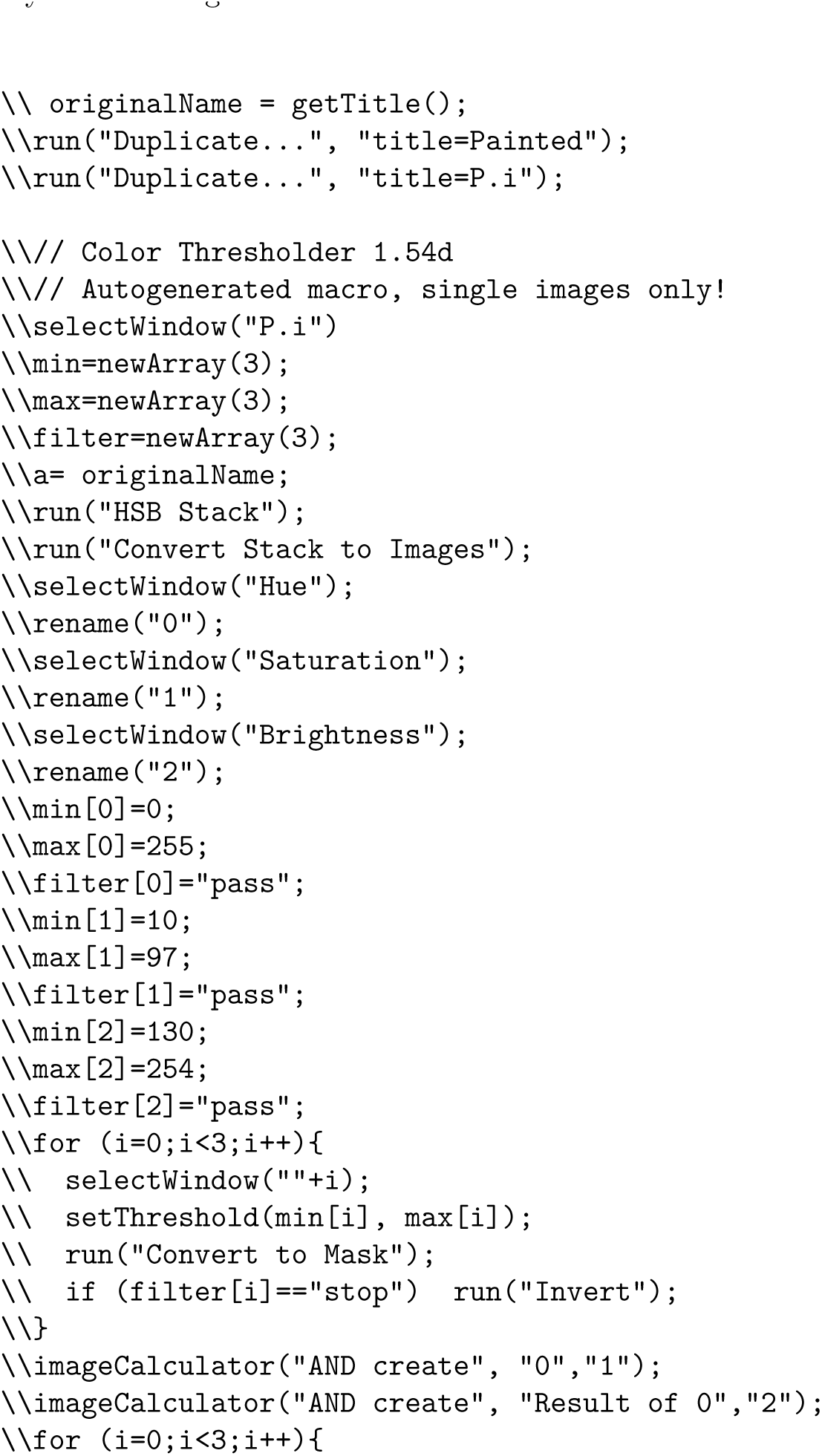
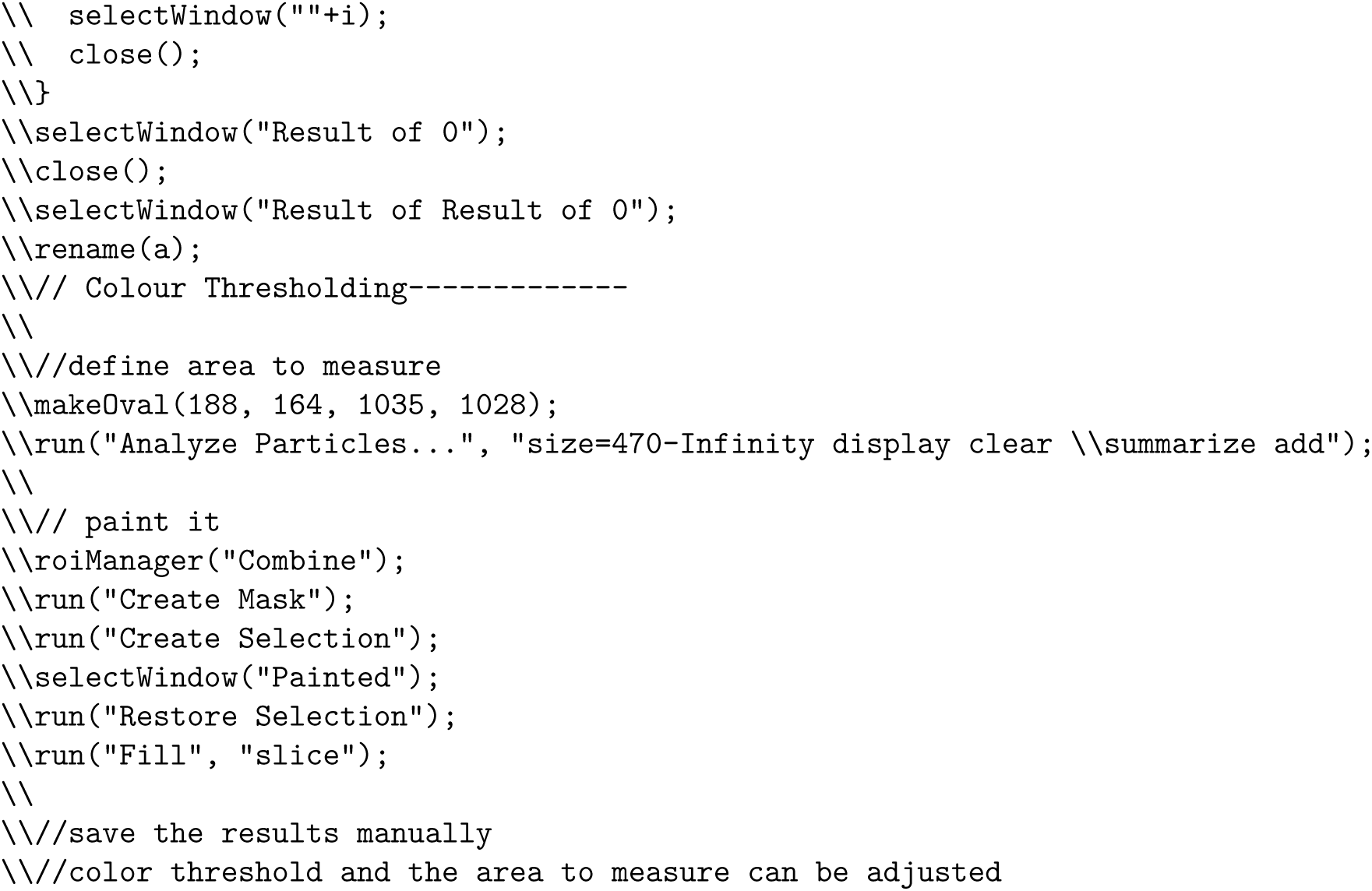

